# System-wide extraction of cis-regulatory rules from sequence-to-function models in human neural development

**DOI:** 10.64898/2026.01.14.699402

**Authors:** Seppe De Winter, Camiel Mannens, Valerie Christiaens, Roel Vandepoel, Lijuan Hu, Sten Linnarsson, Stein Aerts

## Abstract

The genomic *cis*-regulatory code (CRC) underlies spatiotemporal specificity of gene expression. While sequence-to-function (S2F) models can accurately encode the CRC of transcriptional enhancers, decoding these models into human-interpretable rules remains a major challenge. Here we tackle this challenge in human neural development, for which we generate two new single-cell multiome atlases, one from a human embryo and one from neural tube organoids. We use this comparative framework to robustly extract combinations of transcription factor (TF) binding sites that are necessary and sufficient to design enhancers. As such we extract *cis*-regulatory rules for dorsal-ventral progenitors, neural crest, mesenchyme and neurons. To enable this, we develop a new strategy and computational package, called TF-MINDI, to embed, cluster, and annotate candidate TF binding sites, and to extract combinatorial rules for each cell type. We evaluate rule-based models in conjunction with blackbox S2F models through simulations, evolutionary comparisons with zebrafish, topic modeling, and enhancer reporter-assays. Our findings show robust and interpretable rule extraction and constitute a step forward in deciphering, explaining, and formalizing the CRC. TF-MINDI is available at: https://github.com/aertslab/TF-MINDI.

## Introduction

Genomes of multicellular organisms specify all cell types and states^1–3^ through *cis*-regulatory elements, called enhancers^4^. Central to the study of the *cis*-regulatory code (CRC) is the paradigm that enhancers consist of heterotypic clusters of transcription factor binding sites (TFBS)^5,6^ with soft syntactic rules^7^, and that cooperative binding of multiple transcription factors (TFs) underlies cell type-specific enhancer activity^8^, and consequently target gene expression, through co-factor recruitment and enhancer-promoter interactions^5,9^.

Sequence-to-function (S2F) deep learning methods^10–24^ have shown major advances in the ability to model enhancers^7,25^. Several recent S2F applications have illustrated that well-trained S2F models contain accurate representations of the CRC. In other words, the S2F models have *encoded* the CRC. These applications include: (1) the extraction of candidate TFBS through explainability techniques (i.e., contribution scores)^26,27^ and the identification of enriched patterns^26^; (2) the prediction of enhancer activity (e.g., massively parallel reporter assays; MPRA)^14,19,21,22,28,29^; (3) the identification and interpretation of *cis*-regulatory genetic variation^21,23,29–31^; (4) crucially, the successful design of cell type-specific synthetic enhancers^32–35,25,36,37^; and (5) the evolutionary comparison of enhancers and cell types across species^19,38^. Overall, these studies support the CRC paradigm stated above. However, due to the black-box nature of S2F models, it remains challenging to extract human-interpretable rules and generalizable mechanistic models of *cis*-regulation from S2F models.

To address the challenge of CRC extraction, we developed TF-MINDI: **<u>TF m</u>**otif **i**nstance **<u>n</u>**eighborhood **<u>d</u>**ecomposition and **i**nterpretation. TF-MINDI is a computational framework that uses S2F-derived contribution scores to identify TF binding motifs along with their genomic instances. It then embeds these instances in a high-dimensional space, enabling their clustering, annotation, and the modeling of TF binding site co-occurrence patterns. This workflow reveals underlying *cis*-regulatory rules, which can in turn guide the design of novel enhancers for evaluation via S2F models or enhancer reporter assays.

Using TF-MINDI, we extracted the enhancer code of early human development and assessed its accuracy by performing two replicates of the same biological system, generated through independent processes: one from neural tube organoids^39^ and the other from an actual human embryo.

The formation of the neural tube and early neural patterning are highly conserved processes in vertebrates that form the bases for all subsequent neural structures including the central nervous system (from neural tube)^40^, peripheral nervous system (from neural crest and neural tube)^41^ and enteric nervous system (from neural crest)^42^. The neural tube dorsal-ventral axis is established through opposing gradients of SHH (secreted from the notochord and floor plate) causing ventralization, and BMP and Wnt (secreted from the roof plate) causing dorsalization^43^. This gradient in turn drives expression of spatially restricted TFs like *FOXA2*, *NKX2-2*, *PAX5*, *IRX3* and *ZIC1* that establish the dorsal-ventral domains that lead to distinct neuronal subtypes^44^. Likewise, the neural crest is specified at the neural plate border by expression of *SOX10*, *TFAP2A/B*, *PAX3* and *GRHL2*^45–47^. The well-characterized nature of this system combined with the ability to model this early developmental stage *in vitro* makes it an ideal candidate for the development of novel tools for the analysis of the *cis*-regulatory logic. Indeed, 3D organoid protocols have been established for modeling human neural development^39,48–50^, representing a reductionistic biological model system that can be used to interrogate how cell types are encoded in the human genome.

We trained S2F models on chromatin accessibility data of the human neural tube derived from organoids, as well as from an actual human embryo, and using TF-MINDI we extracted concordant enhancer codes underlying early human neural cell types across both systems. We validated the extracted codes using TF ChIP-seq data, synthetic enhancer design and cross-species comparisons of conserved enhancers. Finally, we model TFBS co-occurrences, using topic modeling, and show that its predictive accuracy approaches deep learning S2F models in a cross-species setting.

### An organoid and embryo single-cell multiome atlas of early human neural development

To model early human neural development, we generated two single-cell multiome atlases: one on neural tube organoids^39^ and another on the head of a four weeks post conception human embryo.

For the first atlas, we embedded single cells from two induced pluripotent stem cell (iPSC) lines (Sigma and BJ1) and one embryonic stem cell line (ESC; H9) into poly-ethylene glycol (PEG) based extracellular matrices. The organoids were cultured for 11 days and their medium was supplemented with retinoic acid (RA) and Smoothened agonist (SAG) from day 3 until day 5 (**fig. 1a**). These conditions were previously shown to generate multiple neuronal progenitor and neural crest states^39^. We applied 10x single-cell multiome (combined scATAC-seq and scRNA-seq) on this culture resulting in 22,863 high quality cells (**fig. 1b**) Across the three cell lines similar transcriptomic states were obtained (**fig. S1a-b**) demonstrating the robustness of this organoid system. For the second atlas we performed 10x single-cell multiomics on the head of a four weeks post conception human embryo, resulting in 25,785 high quality cells (**fig. 1c**).

**Figure 1.**
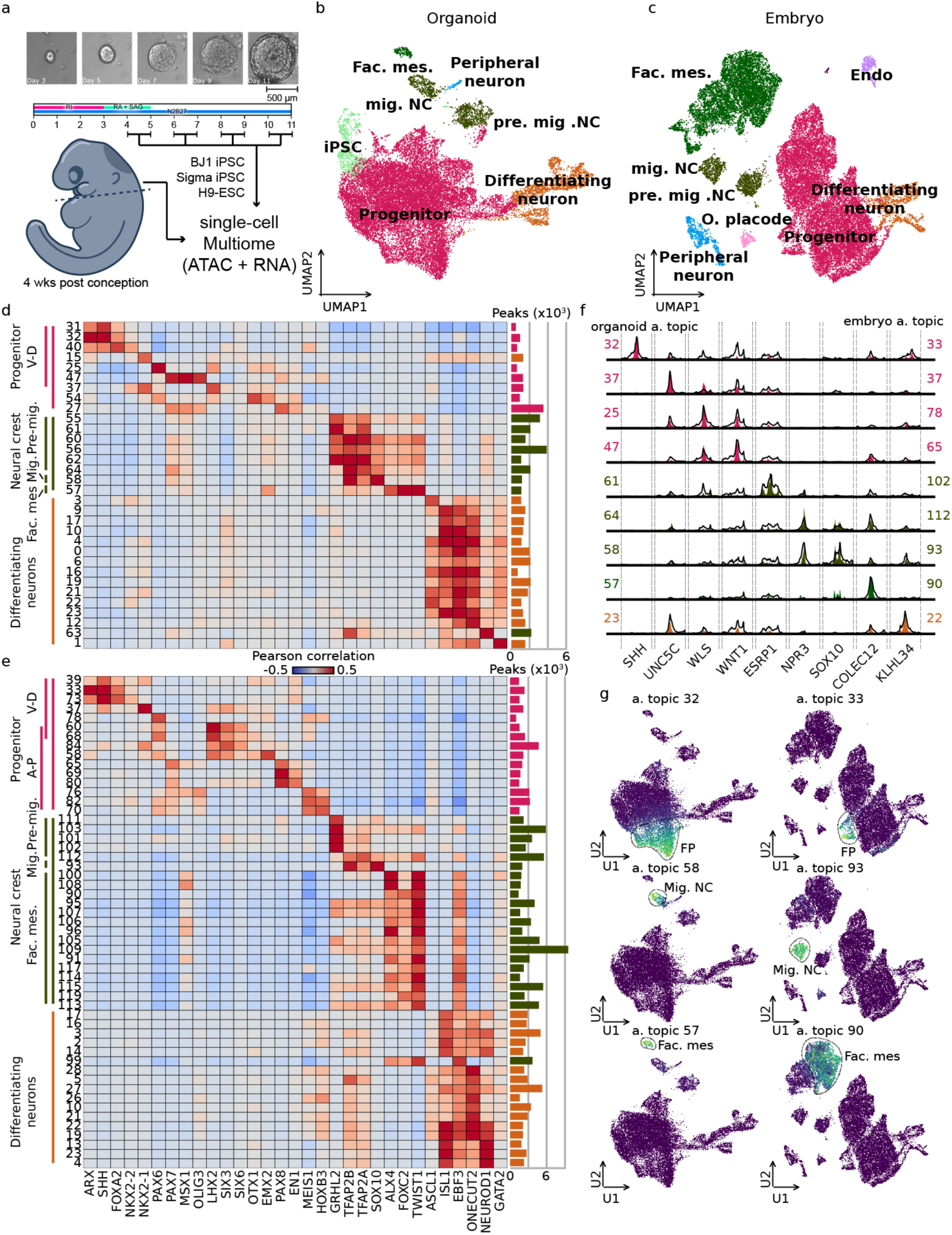
Profiling early human neural development through neural tube organoids and head of four weeks post conception human embryo. **a**, Schematic representation of neural tube organoid protocol and head of a four weeks post conception human embryo on which single-cell multiome was applied. **b-c**, UMAP dimensionality reduction of 22,862 organoid cells (b) and 25,785 embryo cells (c) colored by broad cell type labels. **d-e**, Pearson correlation coefficient between cell topic probabilities and marker gene expression and number of regions per topic for organoid (d) and embryo (e). **f**, Pseudobulk chromatin accessibility profiles per topic for topic regions near marker genes for organoid (filled) and embryo (outline). **g**, UMAP dimensionality reduction colored by example cell topic probabilities in organoid (left) and embryo (right).

We annotated both organoid and embryo cells based on the expression of marker genes (**Table S1**). In both systems we could identify neural progenitors, early differentiating neurons, neural crest (NC), facial mesenchyme and differentiating peripheral neurons (**fig. 1b-c**). Additionally in the organoid only we could identify a pluripotent stem cell population (**fig. 1b**) while perivascular macrophages (PVM) / microglia (MIC) and otic placode (o. placode) cells were only present in the embryo data (**fig. 1c**). For matching cell types, the gene expression profiles are highly correlated across organoid and embryo (**fig. S1c**), with an average Pearson correlation coefficient of 0.89. Annotating cells for cell cycle phases reveals that cells belonging to neuronal clusters are in cell cycle arrest (**fig. S2**), an important hallmark of differentiating neurons^51^, while other cell types are still dividing (**fig. S2**).

Given that the organoid and embryo data represent continuous cell states rather than discrete cell types, we applied topic modeling to the scATAC-seq count matrix, using pycisTopic^52^. This is an unsupervised method to obtain a continuous latent representation of the cell states in the form of chromatin accessibility topics (acc. topics). We performed topic modeling separately for neural progenitors, differentiating neurons and NC (including NC, facial mesenchyme and peripheral neurons), resulting in 75 topics on the organoid data and 120 topics on the embryo data. To annotate topics, we calculated the correlation coefficient between cell-topic probabilities and gene expression of known marker genes (**fig. 1d-g, fig. S3** and **fig. S4**). As such we could identify topics in both organoid and embryo corresponding to neural progenitor dorsal-ventral patterning, marked by the expression of *ARX*, *SHH*, *FOXA2*, *NKX2-2*, *NKX2-1*, *PAX6*, *PAX7*, *MSX1* and *OLIG3* (**fig. 1d-g**, and **Fig. S5**); topics corresponding to anterior-posterior patterning of neural progenitors, only in the embryo data, marked by the expression of *LHX2*, *SIX3*, *SIX6*, *OTX1*, *EMX2*, *PAX8*, *EN1*, *MEIS1*, and *HOXB3* (**fig. 1b**); Topics corresponding to pre- and migratory neural crest, marked by the expression of *GRHL2*, *TFAP2B*, *TFAP2A* and *SOX10* (**fig. 1d-g** and **Fig. S5**); topics corresponding to facial mesenchyme, marked by the expression of *ALX4*, *FOXC2* and *TWIST1*; and topics corresponding to early differentiating neurons, marked by the expression of *ASCL1*, *ISL1*, *EBF3*, *ONECUT2*, *NEUROD1* and *GATA2* (**fig. 1d-g** and **Fig. S5**).

In conclusion, we generated two single cell multiome atlases of early human neural development: one derived from human neural tube organoids and the other from the head of a four week post conception human embryo. Across both datasets, we identify neural progenitors spanning distinct dorsal-ventral identities, pre- and migratory neural crest populations, facial mesenchyme and early differentiating neurons. Gene expression profiles for corresponding cell states were highly concordant between organoid- and embryo-derived cells. Finally, for each cell state we identified cell type-specific chromatin accessibility peaks using topic modeling.

### DeepNeuralTube: sequence-to-function enhancer models of human neural development

To model cell type/state-specific enhancer logic we trained two sequence-to-function (S2F) models to classify genomic regions to *acc. topics* in the organoids (deepNeuralTube°) or embryo (deepNeuralTube^e^) using CREsted^24^ (**fig. 2a**). Both models are able to accurately classify validated enhancers from the Vista enhancer browser^53,54^ as being active in either the neural tube (auROC of 0.75 and 0.80 for resp. deepNeuralTube° and deepNeuralTube^e^ and auPR of respectively 0.21 and 0.29) or facial mesenchyme (auROC of 0.78 and 0.79 for resp. deepNeuralTube° and deepNeuralTube^e^ and auPR of 0.17 for both models; **fig. 2b**). Furthermore, the prediction score of deepNeuralTube° and deepNeuralTube^e^ is highly correlated on these sequences (PCC=0.7; **fig. S6**) showing the robustness of the sequence-to-function modeling approach in this replicate analysis.

**Figure 2.**
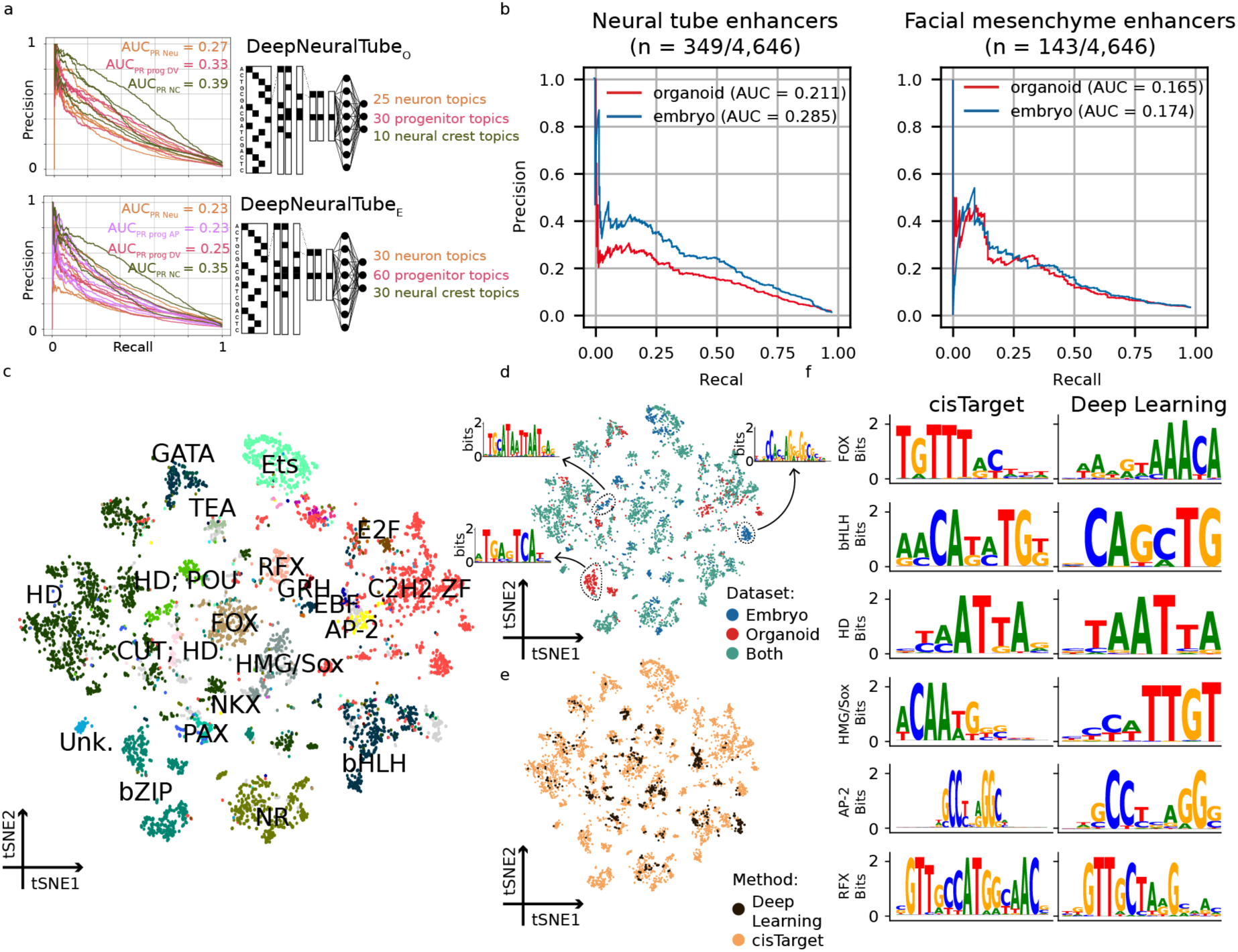
DeepNeuralTube: accurate sequence-to-function models of early human neuro-developmental enhancers. **a**, Precision recall curves for DeepNeuralTube° and DeepNeuralTube^e^ stratified by broad cell type. **b**, precision-recall curves for classifying enhancer active in the neural tube (left) and facial mesenchyme (right) using DeepNeuralTube° and DeepNeuralTube^e^. **c-e**, tSNE dimensionality reduction of 7,630 position weight matrices either from motif enrichment or from DeepNeuralTube contribution scores using TF-MoDISco based on pairwise motif similarity; colored by cluster annotation (**c**), dataset of origin (**d**; in this case only showing motifs from motif enrichment analysis) and whether the motif is obtained from the deep learning models or motif enrichment (**e**). **f**, representative position weight matrix logos from clusters in c.

Next, we explored whether the models learned known TF binding motifs as features in order to make accurate predictions. Therefore, we calculated nucleotide-level contribution scores on 1,000 regions per class, with the highest prediction score, and calculated pairwise motif similarities of contribution score-based motifs (obtained through TF-MoDISco^26^) and known motifs, which are enriched in those same genomic regions. This resulted in clustering of motifs based on TF families (**fig. 2c**). All clusters consist of motifs enriched in both organoid and embryo regions, except for an organoid specific bZIP cluster and an embryo specific C2H2 zinc finger (ZF) cluster (CTCF) and homeodomain; POU (HD; POU) cluster (**fig. 2c-d**). Importantly, most clusters (70 %) contain at least one contribution score-based motif (**fig. 2e-f**).

In conclusion, the two deepNeuralTube models can accurately classify cell type-specific enhancer activity and have learned similar cell type-specific TF binding motifs.

### Concordant enhancer code extraction across human embryo and organoid derived cell states using TF-MINDI

In order to obtain high dimensional embeddings of motifs identified using S2F models, annotate TF binding sites (TFBS) across cell type-specific peaks and model TFBS co-occurrences we developed TF-MINDI. The input to TF-MINDI are contribution scores of regions of interest (**fig. 3a**). Within these contribution scores, TF-MINDI detects seqlets –or motif instances– as spans of nucleotides with high contribution scores. For each such seqlet, pairwise similarity to a set of known motifs is computed, resulting in an embedding per seqlet that is used for clustering and annotation (**fig. 3a**).

**Figure 3.**
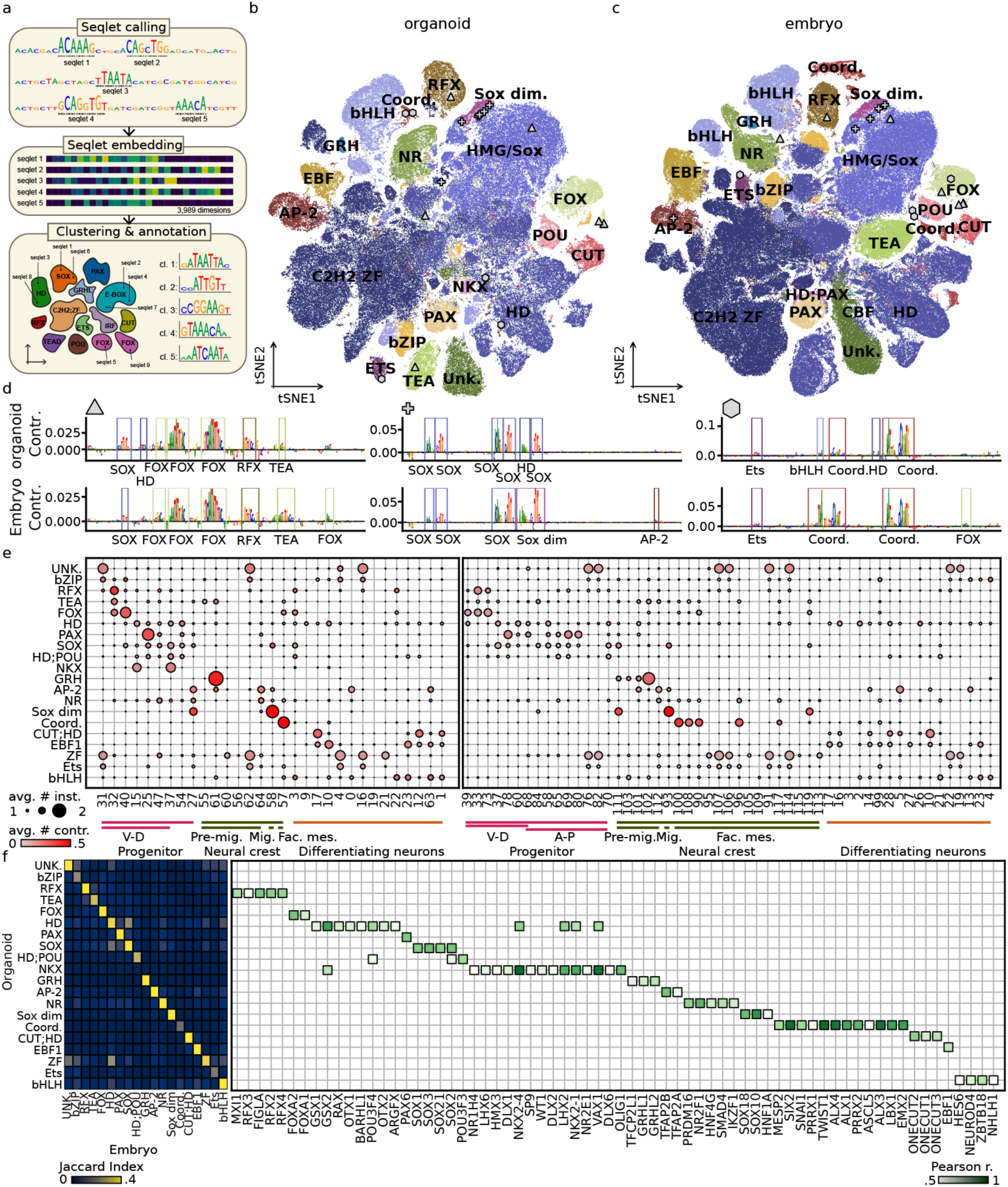
Extraction of transcription factor binding sites through TF-MINDI reveals similar cell type-specific regulators in human neural tube organoid and embryo. **a**, Schematic representation of TF-MINDI workflow. First seqlets are called (i.e., transcription factor (TF) binding instances), for each seqlet the similarity is calculated to a set of known motifs followed by embedding, clustering and annotation of seqlets to TFs and TF-families. **b-c**, tSNE dimensionality reduction of 445,039 organoid seqlets (**b**) and 668,928 embryo seqlets (**c**) colored based on TF-family annotation. **d,** Contribution score for representative floor plate (chr5:83085049-83085549; hg38), migratory neural crest (chr16:972507-973007; hg38) and facial mesenchyme (chr6:165721333-165721833; hg38) peaks using DeepNeuralTube° (top) and DeepNeuralTube^e^ (bottom) annotated automatically using TF-MINDI. Symbols indicate seqlet locations in panel c and d. **e**, organoid (left) and embryo (right) code-table showing average number of TF binding instances (dot size) and average contribution score (color) per region and cell type. **f**, Jaccard index (left) of the genomic overlap of instances from organoid and embryo TF-MINDI and Pearson correlation coëfficient (right) between average gene expression profile per cell type and average number of instances per region and cell type for TFs annotated to each TF family (only showing values above 0.5) for organoids.

We applied TF-MINDI to contribution scores on 1,000 regions per class of both DeepNeuralTube° and DeepNeuralTube^e^, resulting in resp. 446,038 and 668,928 seqlets. These seqlets were embedded by computing their similarity to 3,989 known-motifs. Based on this embedding, all seqlets were projected into two-dimensional space using principal component analysis (PCA) followed by t-distributed stochastic neighbor embedding (tSNE), resulting in the co-clustering of similar seqlets. As such, we identified seqlets annotated to a total of 19 major TF-families in both organoid and embryo (**fig. 3b-c**). Similar proportions of seqlets are annotated to each family across models (Pearson correlation of 0.98). This analysis results in automated annotations of TFBS instances across cell type-specific peaks (**fig. 3b-d**). Note that in both systems we identified a cluster representing a homeobox and E-box dimer motif (**fig. 3b-d**). This dimer motif, named “coordinator”, was previously shown to be an important regulator in facial mesenchyme^55^ and is bound by TWIST1 and a variety of homeodomain factors including ALX4^56^. Indeed, we found instances of this motif in facial mesenchyme-specific regions (**fig. 3d**).

The automated TFBS instance annotations allow for the quantification of TFBS instance occurrence across cell type-specific peaks, resulting in an enhancer-code table for organoid and embryo (**fig. 3e**). This code table reveals cell type-specific regulators. For example, RFX, FOX and TEAD in floor plate-specific regions (*acc. topic* 32 and 33 in resp. organoid and embryo; (**fig. 3e**)); Grainyhead (GRH) and TEAD in pre-migratory/epithelial NC specific regions (*acc. topic* 61 and 102 in resp. organoid and embryo; (**fig. 3e**)); SOX homodimer, AP-2 and nuclear receptor (NR) in migratory NC specific regions (*acc. topic* 58 and 93 in resp. organoid and embryo; (**fig. 3e**)); Coordinator in facial mesenchyme specific regions (*acc. topic* 57 and 100/108/90 in resp. organoid and embryo; (**fig. 3e**)); and Onecut (CUT;HD), EBF1 and bHLH instances in differentiating neuron specific regions (**fig. 3e**). Not only similar TF families are recovered across cell type-specific regions in organoid versus embryo but there is also a strong overlap (average Jaccard index of 41 %) of their genomic location (**fig. 3f**).

Next, we calculated the Pearson correlation coefficient between average TF expression profile per cell type and average number of instances across cell type-specific regions, to link TFBS instances to TFs (**fig. 3f**). As such, for each TF family, multiple paralog TFs could be linked. For example, both *FOXA1* and *FOXA2* for the FOX family (**fig. 3f**).

Finally, to evaluate how the enhancer code is conserved over time and to assess if motif spaces can be integrated across experiments, we applied TF-MINDI to a published dataset of the developing human brain generated in the same lab at a later developmental timepoint (6-14 weeks post conception)^57^. We found that most TF families were represented at both early and later timepoints and integrated well without performing batch effect correction (**fig. S7a-b**), but that some transcription factors had well-defined developmental timings. For instance, NFI factors (part of the SMAD/NFI family) are only found in the first-trimester dataset, while Grainyhead is only found in the neural tube datasets (**fig. S7c-d**; Supplementary note 1).

In conclusion, TF-MINDI automatically extracts, embeds, and clusters TFBS instances underlying cell type-specific regions of human neural development. Through this analysis, we obtained concordant enhancer codes across organoid- and embryo-derived cell types.

### Validation of TF-MINDI transcription factor binding instances through ChIP-seq in facial mesenchyme and peripheral blood mononuclear cells

To validate TFBS instances predicted using TF-MINDI, we made use of TWIST1 chromatin immunoprecipitation and sequencing (ChIP-seq) data in facial mesenchyme cells. Seqlets annotated as Coordinator are enriched for TWIST1 ChIP-seq signal (**fig. 4a-b**), and TF-MINDI instances result in a higher auROC compared to motif enrichment analysis using pycisTarget^52^ (**fig. 4c**). Furthermore, TF-MINDI coordinator seqlets recover a large fraction of the genome-wide TWIST1 ChIP-seq peaks (2,483 peaks are recovered from the top 10,000 ChIP-seq peaks; **fig. 4d**).

**Figure 4.**
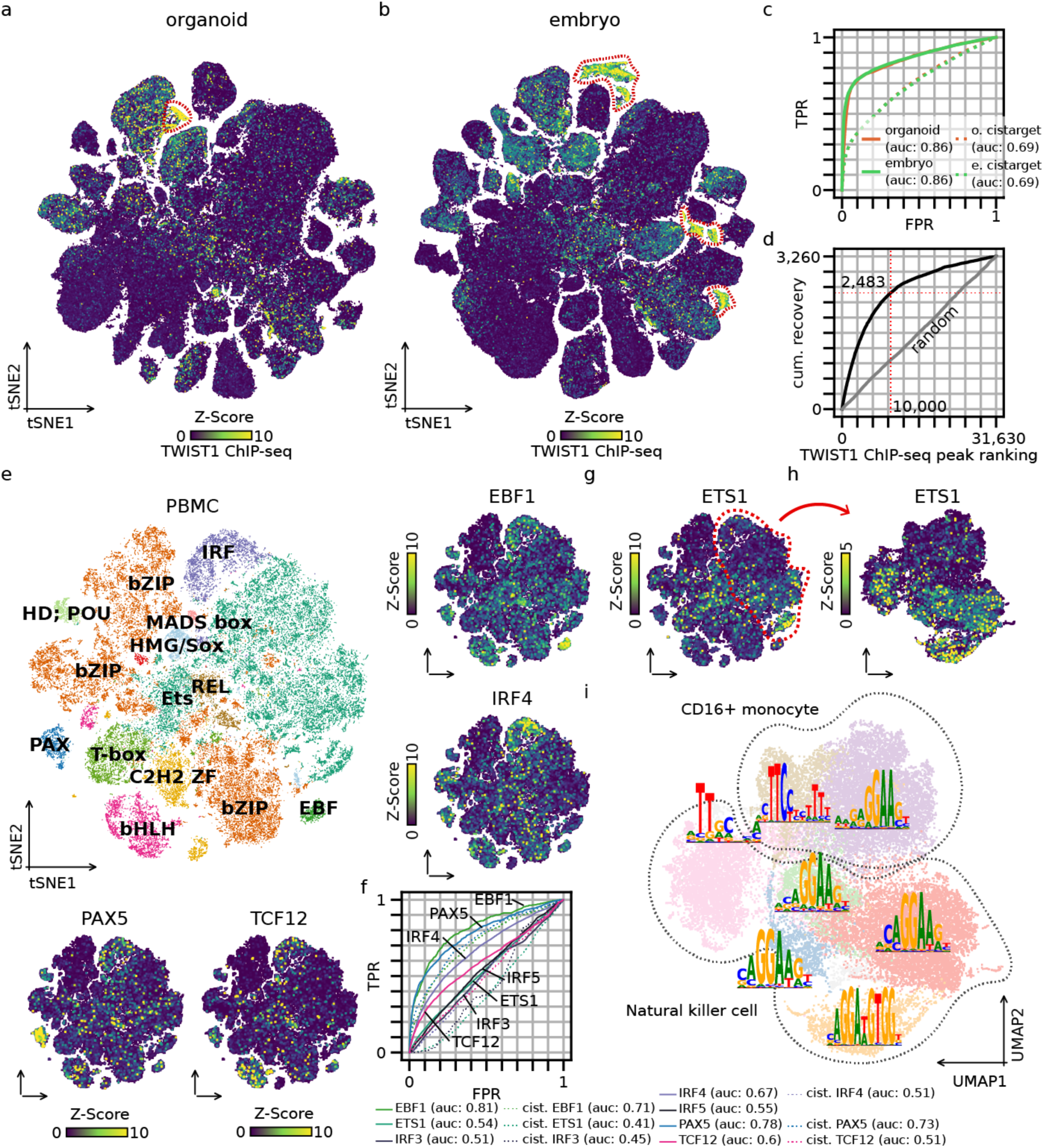
Enrichment of ChIP-seq signal supports TF-MINDI predicted transcription factor binding sites. **a-b**, tSNE dimensionality reduction of 445,039 organoid seqlets (**a**) and 668,928 embryo seqlets (**b**) colored by TWIST1 ChIP-seq signal in facial mesenchyme cells; location of coordinator seqlets is marked by red dotted line **c**, ROC curve showing classification of TWIST1 ChIP-seq signal (across all seqlets) using TF-MINDI coordinator seqlets or using TWIST1 motif enrichment (cistarget). **d**, Recovery curve showing genome-wide recovery of TWIST1 ChIP-seq peaks using TF-MINDI coordinator seqlets; the dotted line marks the top 10,000 peaks. **e**, tSNE dimensionality reduction of 74,204 seqlets from deepPBMC colored by TF-family and EBF1, IRF4, PAX5 and TCF12 ChIP-seq signal in the GM12878 cell line. **f**, ROC curve showing classification of TF ChIP-seq signal (across all seqlets) using TF-MINDI seqlets of the corresponding TF-family or using motif enrichment (cist.). **g-h**, tSNE dimensionality reduction of 74,204 seqlets from deepPBMC (**g**) and UMAP dimensionality reduction of a subset 23,959 seqlets annotated to the Ets family colored by ETS1 ChIP-seq signal in the GM12878 cell line. **i**, UMAP dimensionality reduction of a subset 23,959 seqlets annotated to the Ets family from deepPBMC colored by cluster annotation and position weight matrix logos per cluster. Cell type-specific peaks from which the seqlets originate are indicated with a dashed line.

To further extend this validation in other biological systems, we applied TF-MINDI to cell type-specific regions from peripheral blood mononuclear cells, (PBMC) using the deepPBMC S2F model^24^. TF-MINDI identified 74,204 seqlets that were annotated to 13 TF-families (**fig. 4e**). To validate the identified TFBS instances we made use of EBF1, IRF4, PAX5 and TCF12 ChIP-seq data in the B-cell lymphoblastoid cell line GM12878. Indeed, we observe specific enrichment of ChIP-seq signal in seqlets annotated to the corresponding TF families (**fig. 4e-f**). Also in this dataset, TF-MINDI instances result in a higher auROC compared to motif enrichment analysis using pycisTarget^52^ (**fig. 4f**).

Depending on the Leiden clustering resolution, TF-MINDI yields multiple clusters for each TF-family (28 clusters for 13 families in this PBMC example). For example, seqlets annotated as ETS show particularly high heterogeneity (**fig. 4e**). To test if this variation is biologically meaningful, we subclustered ETS instances and evaluated ETS1 ChIP-seq signal (**fig. 4g-h**). In this case, we observed that only a subset of ETS seqlets showed a high ChIP-seq signal (**fig. 4h**). Generating position weight matrix (PWM) logos for each of the Ets sub-clusters reveals that instances with higher ChIP-seq signal tend to be of the form CMGGAAG while those with lower signal tend to be of the form GRGGAAG (**fig. 4i**). The former instances predominantly originate from natural killer cell peaks, while the latter predominantly originate from CD16+ monocyte peaks (**fig. 4h**). This suggests that the heterogeneity observed for ETS instances is biologically meaningful. Indeed, the consensus TFBS sequence of ETS1 is CMGGAAG^58^.

In conclusion, TF-MINDI clusters represent biologically relevant TFBS instances. These clusters do not only represent instances of different TF-families, they can also represent differences in TF-affinity.

### Validation of the TF-MINDI extracted facial mesenchyme enhancer code through synthetic enhancer design

If the extracted enhancer code rules are sufficient, they should, in principle, enable the design of novel enhancers that score highly according to DeepNeuralTube and that, when tested using enhancer reporter assays, exhibit cell type-specific enhancer activity.

To test whether using TF-MINDI we can extract such rules, we focused on facial mesenchyme cells, which are relatively easy to culture in the laboratory. For this purpose, we applied TF-MINDI to all 7,296 organoid facial mesenchyme genomic regions (i.e. *acc. topic* 57 regions). TF-MINDI identified 40,249 seqlets annotated to 11 TF-families (**fig. 5a**). A subset of seqlet clusters represent a homeobox and E-box dimer motif called the coordinator^55,56^. We were able to increase the number of instances annotated as this dimer motif using the TF-MINDI dimer detection module, that we developed for this purpose (**fig. 5b**). This method works by centering contribution scores based on a query seqlet cluster, aggregating the signal across all instances and detecting additional peaks at a fixed distance to the query instances (**fig. 5b**). Using this approach we could detect 4,999 coordinator instances across cell type-specific peaks.

**Figure 5.**
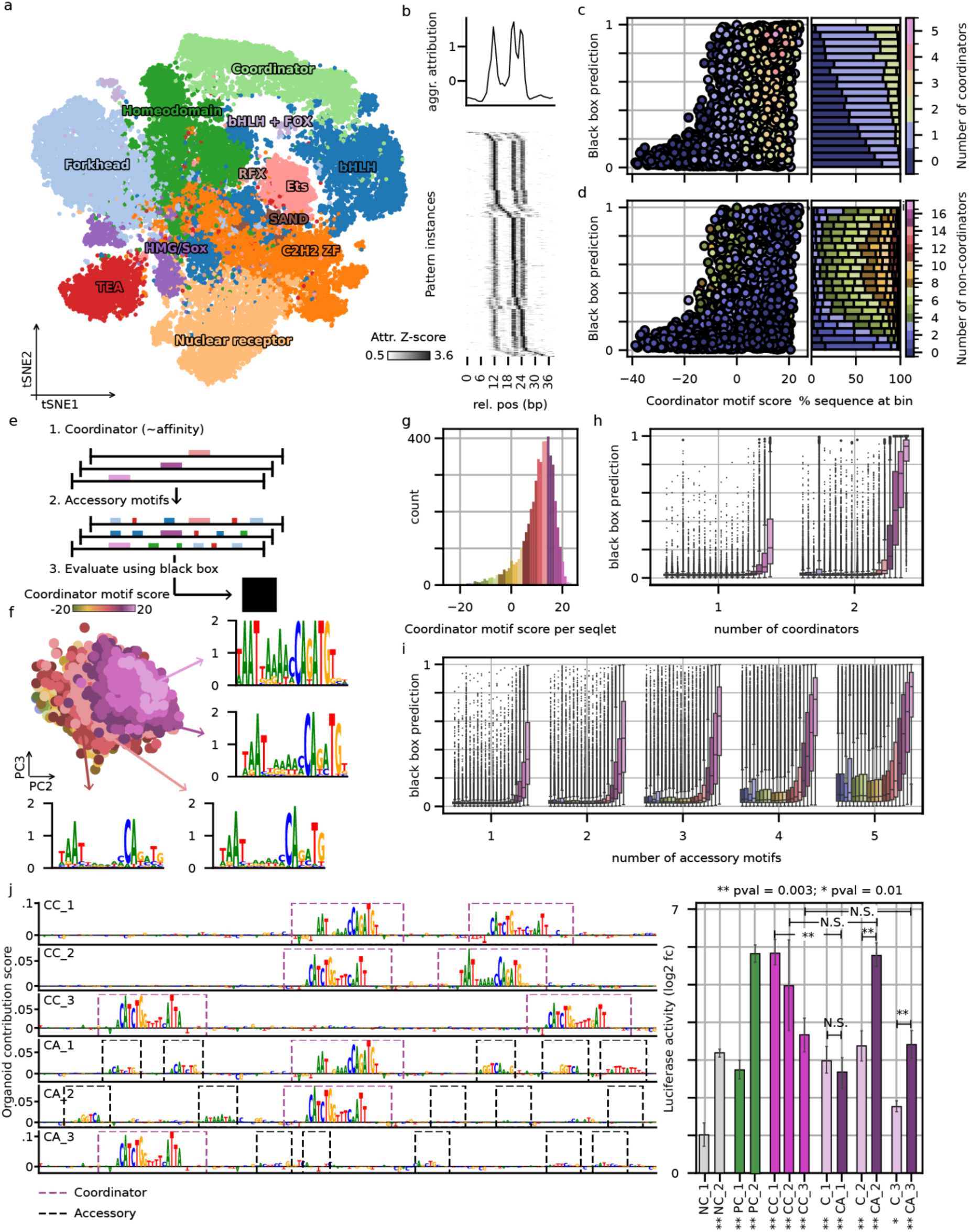
Extraction of the facial mesenchyme enhancer code using TF-MINDI, validated via synthetic enhancer design. **a**, tSNE dimensionality reduction of 40,249 organoid facial mesenchyme seqlets colored by TF-family. **b**, illustration of transcription factor binding site dimer detection showing contribution score profile across aligned coordinator seqlets; top plot shows average profile across seqlets. **c-d**, Scatter plot of facial mesenchyme regions (n=7,296) relating coordinator position weight matrix (PWM) score to DeepNeuralTube° (black box) prediction score colored based on number of coordinator instances (c) or non-coordinator instances (d). **e**, Illustration of enhancer design strategy: first up to two coordinator instances of varying affinity are implanted in random DNA sequences followed by up to five accessory instances. **f**, PCA of the subset 4,999 coordinator seqlets colored by coordinator PWM score binned according to distribution in panel g. **g**, Distribution of coordinator PWM score across coordinator seqlets. **h-i**, DeepNeuralTube° (black box) prediction score after inserting one or two coordinator instances of varying PWM scores (color) into 10,000 random DNA sequence (**h**), and after adding up to five accessor instances to sequences containing a single coordinator instance at varying affinity (color; **i**). **j**, Left: organoid contribution scores for experimentally tested synthetic enhancers, sampled from CC (two high-affinity coordinator instances) and CA (one high-affinity coordinator instance plus five accessory instances). Right: luciferase reporter assay in facial mesenchyme cells (n=6 sequences per condition; 3 technical replicates measured across 2 biological replicates). NC: negative control (genomic region with low DeepNeuralTube prediction score); PC: positive control (genomic region with high DeepNeuralTube prediction score). Two-sided Mann–Whitney U test was used to test for the difference in mean. P-values are corrected using the Benjamini-Hochberg procedure. Stars under x tick labels represent test compared to NC_1.

Given the importance of the coordinator motif for facial mesenchyme enhancers^55,56^ we assessed how its affinity, quantified using the motif score of a known coordinator motif, across facial mesenchyme peaks relates to the DeepNeuralTube° facial mesenchyme prediction score (**fig. 5c-d**). Regions with a high DeepNeuralTube° facial mesenchyme prediction score tend to have a high coordinator affinity. However, not all regions with a high affinity for the coordinator also have high prediction scores (**fig. 5c-d**). Quantifying the number of coordinator (**fig. 5c**) and non-coordinator (**fig. 5d**) instances per region reveals that most regions with a high facial mesenchyme prediction score have one or two coordinator instances but also several (up to 10) non-coordinator instances (**fig. 5c-d**). Note that some regions in the genome have multiple coordinator copies and no additional instances while having a high facial mesenchyme predictions score.

From this we hypothesize that facial mesenchyme-specific enhancers are encoded using a code that consists of two levels. The first being the presence of one or more copies of coordinator and the second being the presence of additional instances of other TF families.

To test this hypothesis, we designed synthetic enhancers by first implanting one or two coordinator instances at random positions within a 160 bp window in 10,000 random DNA sequences following the local GC content of scATAC-seq peaks (**fig. 5e**). Next, we added up to five additional TFBS instances in the same 160 bp window, ensuring a minimal distance of 3 bp between binding sites (**fig. 5e**). These instances were sampled from the TF-MINDI clusters, taking the top 5 % most frequently occurring kmers per cluster. We also assessed the effect of coordinator affinity. Subclustering coordinator instances reveal an affinity gradient (**fig. 5f-g**) and generating PWM logos at different affinity bins (**fig. 5g**) highlights the importance of adenine nucleotides in between the homeobox and E-box sites (**fig. 5g**). Accordingly, for the enhancer design experiments we implanted coordinator instances sampled from different affinity bins.

In line with the quantification of coordinator instances in genomic regions (**fig. 5c**), implanting two high-affinity coordinator instances is sufficient to generate synthetic enhancers according to both DeepNeuralTube° (**fig. 5h**) and DeepNeuralTube^e^ (**fig. S8**). However, implanting only a single instance is not sufficient (**fig. 5h**; **fig. S8**). Starting from genomic regions in which one high-affinity coordinator instance has already been implanted, the additional implantation of four or five TFBS is sufficient (**fig. 5i**; **fig. S8**).

To validate these rules experimentally, we implanted one high-affinity coordinator instance in three random DNA sequences and either implanted an additional high-affinity coordinator instance (CC_1, CC_2, and CC_3) or five additional non-coordinator instances (CA_1, CA_2, and CA_3) using the same design rules as described above. Next we measured enhancer activity in *in vitro* differentiated facial mesenchyme cells using a luciferase reporter assay, together with two negative controls (i.e., genomic regions with low prediction score for both DeepNeuralTube models; NC_1 and NC_2) and two positive controls (i.e., genomic regions with high prediction score for both DeepNeuralTube models; PC_1 and PC_2). All DNA sequences containing two high-affinity coordinator instances exhibit strong enhancer activity (**fig. 5j**), whereas this activity is reduced when only a single instance is present (**fig. 5j**). The additional implantation of five non-coordinator TFBS into DNA sequences that already contained a single high-affinity coordinator instance increases their enhancer activity to levels close to those of sequences with two high-affinity coordinator instances in two out of three cases (**fig. 5j**). Furthermore, the luciferase activity is correlated (PCC 0.67) to the DeepNeuralTube prediction score (**fig. S9**). Surprisingly, one negative control DNA sequence (NC_2) had relatively high enhancer activity (**fig. 5j**). Evaluating this sequence using DeepNeuralTube° reveals that it has a high prediction score for migratory neural crest cells, containing a SOX dimer, bHLH and AP-2 motif (**Fig. S11**). Its enhancer activity might thus be the result of migratory neural crest cell contamination in the facial mesenchyme culture. Finally, to assess the cell type-specificity of these synthetic enhancers we performed an additional luciferase reporter assay in a melanoma cell line (MM001). In this cell line, none of the enhancers are active (**fig. S10**).

To conclude, using TF-MINDI we extracted the facial mesenchyme enhancer code. We show, both computationally and experimentally, that the presence of either two high-affinity coordinator instances or a single high-affinity coordinator instance with additionally five accessory TFBS is sufficient to generate synthetic enhancers.

### Cross-species enhancer-code integration and modeling of transcription factor binding site co-occurrences

Cell type-specific enhancer activity is not only encoded through the presence of individual TFBS (and their copy number and affinity) but also by the co-occurrence of multiple instances. To model this aspect of the enhancer code, we employ the probabilistic framework of topic modeling^59^. For this purpose, we generated a count matrix containing the number of TFBS instances across genomic regions (**fig. 6a**). The features of this matrix correspond to the clusters obtained by TF-MINDI, intrinsically modeling variations in TF-motif instances, for example reflecting differences in TF affinity. The topic modeling procedure iteratively optimises two probability distributions: (1) the probability of a TF-MINDI cluster belonging to a topic and (2) the contribution of a topic to a genomic region (**fig. 6a**). We refer to these topics as pattern-topics (*pat. topics*) to distinguish them from *acc. topics*.

**Figure 6.**
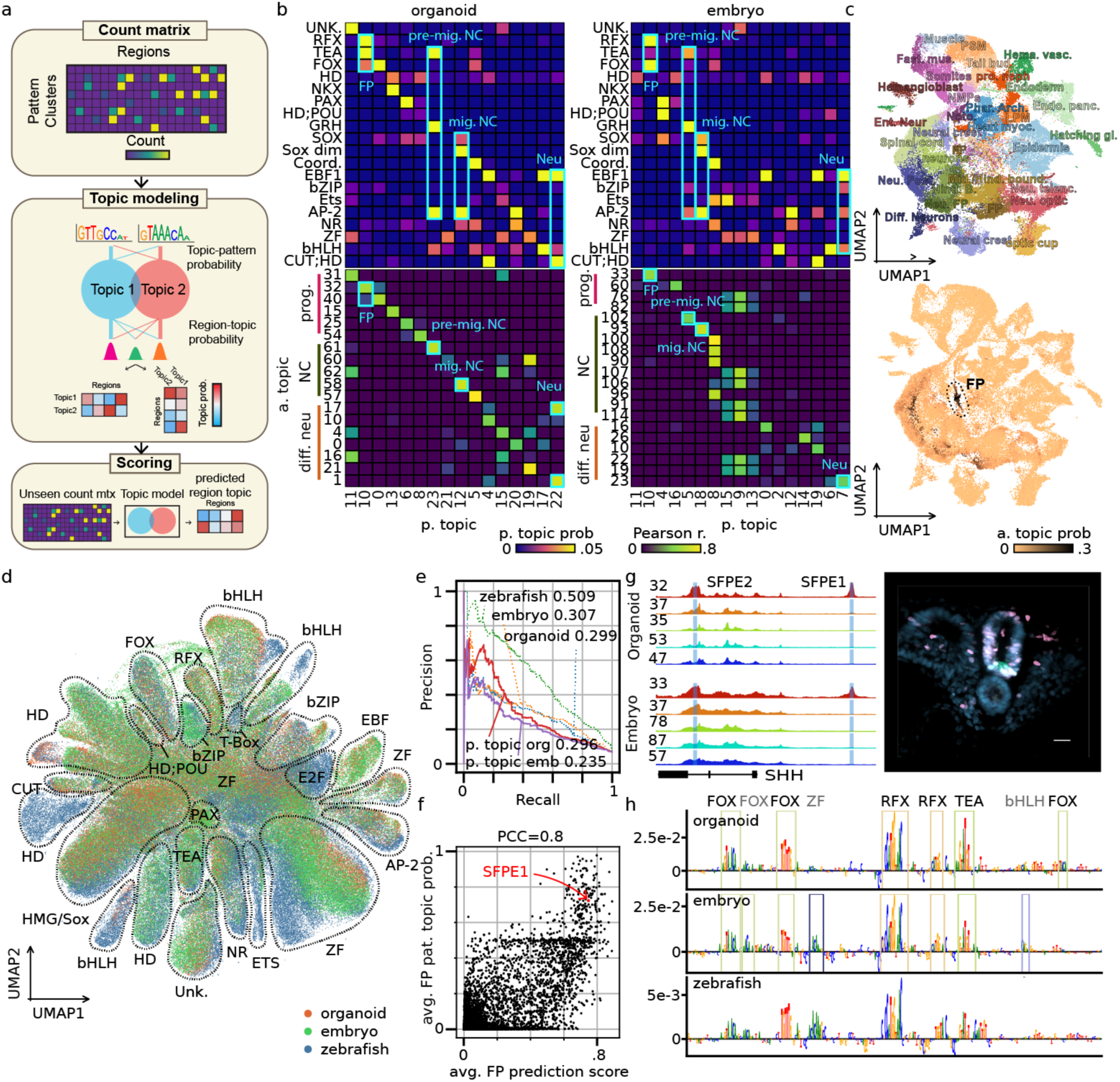
Quantitative enhancer code model capturing transcription factor binding site instance co-occurrence patterns using TF-MINDI topic modeling, validated using cross species analysis. **a**, schematic representation of TF-MINDI topic modeling: a count matrix of regions x TF-MINDI clusters is generated next topic modeling is performed simultaneously optimizing region-topic and pattern-cluster-topic probabilities. The resulting topic model can be used to score unseen count matrices. **b**, Top: pattern-topic probabilities for organoid and embryo, averaged over clusters for each TF family. Bottom: Pearson correlation of region-pattern-topic probabilities with region-accessibility-topic probabilities. **c**, UMAP dimensionality reduction of 104,129 zebrafish cells colored by cell type label (top) and cell topic probability of a floor plate acc. topic (bottom). **d**, UMAP dimensionality reduction of 1,707,039 seqlets from organoid, embryo and zebrafish data colored by dataset of origin; TF-familes are indicated. **e**, precision-recall curve showing classification of zebrafish floor plate topic by the zebrafish model, deepNeuralTube°, deepNeuralTube^e^ and TF-MINDI pattern topic models for organoid and embryo; area under the precision recall curves are indicated. **f**, Scatter plot of 16,293 human peaks for average floor plate (FP) prediction score of DeepNeuralTube° (acc. topic 32) and DeepNeuralTube^e^ (acc. topic 33) and average FP pattern topic probability (pat. Topic 10). SFPE1 is indicated. **g**, Left: chromatin accessibility in organoid (top) and embryo (bottom) across dorsal-ventral topics for locus chr7:155,799,664-155,827,483 (hg38) with SFPE2 and SFPE1 highlighted. Right: chicken embryo enhancer reporter assay: DAPI (blue), electroporation control (PINK), GFP driven by SFPE1 (green). The scale bar represents 500μm. **h**, Organoid (top), embryo (middle) and zebrafish (bottom) contribution score of SFPE1 and TF-MINDI annotations for organoid and embryo.

We summarised the pattern-topic probabilities at the TF-family level by averaging across clusters annotated to the same TF-family, and related pattern-topics to accessibility-topics by calculating the correlation coefficient between pattern-topic and accessibility-region-topic probabilities (**fig. 6b**). For both organoid and embryo, this revealed a *pat. topic* with high probability for RFX, TEAD and FOX TFBS (*pat. topic* 10 in both systems; **fig. 6b**), that correlates with a floor plate *acc. topic* (*acc. topic* 32 and 33 in resp. organoid and embryo; **fig. 6b**); a *pat. topic* with high probability for GRH, TEAD and AP-2 TFBS (*pat. topic* 23 and 5 in resp. organoid and embryo; **fig. 6b**), that correlates with a pre-migratory NC *acc. topic* (*acc. topic* 61 and 102 in resp. organoid and embryo; **fig. 6b**); a *pat. topic* with high probabilities for SOX-dimer, SOX monomer and AP-2 TFBS (*pat. topic* 12 and 18 in resp. organoid and embryo; **fig. 6b**), that correlates with a migratory NC *acc. topic* (*acc. topic* 58 and 93 in resp. organoid and embryo; **fig. 6b**); and a *pat. topic* with high probability for EBF1, bHLH and CUT;HD (only in organoid) TFBS (*pat. topic* 22 and 7 in resp. organoid and embryo; **fig. 6b**), that correlates with a differentiating neuron topic (*acc. topic* 1 and 23 in resp. organoid and embryo; **fig. 6b**).

To further validate the pattern-topic enhancer code models, we assessed their evolutionary conservation by scoring chromatin accessibility peaks derived from a zebrafish developmental atlas^60^. For this purpose, we reanalysed the scATAC-seq side of a publicly available single-cell multiomics dataset of the developing zebrafish embryo, comprising six developmental stages and 30 annotated cell types^60^. After quality control, we retained 104,129 cells, which were used for topic modeling with a model comprising 100 topics. UMAP dimensionality reduction based on cell topic probabilities clustered the data into the previously annotated cell types, from which we identified an *acc. topic* that is specific to floor plate cells (**fig. 6c**). Next, we trained a classification S2F model, classifying zebrafish genomic regions into the 100 *acc. topics* and applied TF-MINDI resulting in 593,072 seqlets, which we integrated with the seqlets obtained from organoid and embryo (**fig. 6d**).

This integration (**fig. 6d**) results in a shared cross-species TF-MINDI cluster annotation, which enables the scoring of zebrafish chromatin accessibility peaks using the human pattern-topic models. We tested whether the floor plate pattern-topic model (*pat. topic* 10 in both organoid and embryo; **fig. 6b**) could be used to classify floor plate-specific zebrafish chromatin accessibility peaks (**fig. 6e**). Surprisingly, this achieved a precision and recall almost on par with the DeepNeuralTube models (auPR of 0.235 vs 0.299 for organoid, and 0.235 vs 0.307 for embryo), suggesting that a topic model that captures TFBS co-occurrences is sufficient to model cell type-specific enhancers with performance comparable to deep learning models, for this cell type. Furthermore, the prediction score from both DeepNeuralTube models for the floor plate class are highly correlated with region-pattern-topic probabilities of both organoid and embryo floor plate *pat. topics* (*pat. topic* 10 in both) on cell type-specific peaks (PCC=0.8; **fig. 6f**).

As an illustrative example we focus on the *SHH* floor plate enhancer 1 (SFPE1), located upstream of *SHH* (**fig. 6g**), which has been shown to be specifically active in floor plate cells^61^. Its sequence and activity are highly conserved: when we test this enhancer in chicken embryos, SFPE1 drives enhancer activity specifically in the ventral neural tube (**fig. 6g**). This enhancer ranks highly according to both the organoid and embryo floor plate pattern-topic models (**fig. 6f**). Furthermore, both DeepNeuralTube models as well as the zebrafish model produce similar nucleotide explanations, identifying multiple FOX and RFX binding sites and a single TEAD binding site (**fig. 6h**).

In conclusion, modeling cell type-specific enhancers using TFBS co-occurrences through topic modeling yields promising insights into the combinatorial logic of enhancers and achieves predictive performance comparable to deep learning models.

## Discussion

We set out to define the *cis*-regulatory rules of genomic enhancers underlying early human neural development. To this end, we generated two single-cell multiome atlases of human neural tube development derived through two distinct routes: organoid cultures and actual human development and trained sequence-to-function (S2F) deep learning models predicting cell type-specific chromatin accessibility across cell states representing dorsal-ventral progenitors, neural crest and neurons.

To enable the extraction of *cis*-regulatory rules from these models we developed TF-MINDI. This new computational framework facilitates: (1) the *de novo* discovery of TF binding motifs, (2) genome-wide identification of TF binding motif *instances*, (3) high-dimensional embedding, clustering, and annotation of these instances, and (4) modeling TF binding sites (TFBS) co-occurrences. By applying TF-MINDI to two independent replicates (organoids and human embryonic development) of the same biological system we show how it reproducibly extracts *cis*-regulatory rules at a system-wide level.

Using TF-MINDI, we annotated more than 400,000 TFBS instances across cell type-specific chromatin accessibility peaks. A large proportion of these instances (on average ∼40%) were identified in both the organoid and embryo datasets and annotated to the same TF-family, and are therefore considered highly confident. We further validated these annotations using TF ChIP-seq data, further increasing confidence. These annotations were then used to quantify which combination of TFBS constitute cell type-specific enhancers and were linked to TFs based on gene expression, often resulting in multiple paralog candidates for a given TF-family. Importantly, the different instances identified using TF-MINDI do not only represent different TF-families but also differences in affinity for a given TF(-family). For facial mesenchyme cells, we validate the sufficiency of the extracted TFBS by implanting them in random DNA sequences, within a 160 bp window, and showing that this generates cell type-specific enhancers. Finally, using topic modeling, we model TFBS co-occurrences and show, for floor plate cells, that these probabilistic rule-based models can approach the accuracy of deep learning S2F models in a cross-species setting. We believe that the resulting annotation of part of the *cis-*regulatory genome will aid with future endeavors of explaining non-coding variation and that white-box modeling will result in mechanistic insights into *cis*-regulation.

TF-MoDISco^26^ is a widely used computational method for identifying enriched motifs from contribution scores generated through S2F models and it has been successfully applied to uncover sequence determinants in a variety of biological processes^19,20,23,28,38,57,62–74^. With TF-MINDI we broaden the scope of pattern, instance, and rule discovery. A key innovation is to work with a high-dimensional representation (or ‘embedding’) of each individual seqlet, that we construct through a cost-efficient similarity score with a set of known position weight matrices. This enables the genome-wide analysis of the actual TF motif instances, without the need to perform post-hoc scanning of contribution scores. The implementation furthermore makes use of the AnnData^75^ data structure and best practices from the scverse^76^ project for easy interoperability and providing familiar application programming interfaces (APIs).

The *cis*-regulatory rules examined in this work represent only a subset of the full rules underlying the *cis*-regulatory code (CRC). We particularly focused on TFBS and combinations in cell type-specific enhancers, but we envision that in the future additional CRC aspects can be investigated. A first aspect, which is a limitation for current sequence-to-function models, is the modeling of transcriptional repression whereas current sequence-to-function model approaches predominantly identify activator TFs. Transcriptional repression is important for cell type-specific enhancer activity, particularly during neural tube patterning^77^. For example, the GLI TF family, which plays an important role in dorsal-ventral patterning, and has been shown to possess both activator and repressor functions^78^, was not detected by either of the two DeepNeuralTube models. A second aspect is modeling gene expression. To achieve that, enhancers need to be associated with target genes, which is still a major challenge in the field^79–82^. Whereas in this study we focused on enhancer decoding using “local” S2F models trained on enhancer sequences, the link to target gene expression will likely require S2F models that predict gene expression, trained on larger input sequences^21,23,29^. However, whereas these large-input models perform well on the prediction of gene expression for housekeeping genes, or genes expressed in “broad” cell types, cell type-specific enhancers are often missed^24^. Bridging the strengths of the two approaches (local enhancer modeling and large scale gene locus modeling), will likely yield improved CRC-models in the future. A third aspect is that enhancers are nodes in a gene regulatory network (GRN) underlying dynamic cell states^4^. A CRC should ultimately yield the GRN of a biological system, encompassing gene expression dynamics, positive and negative feedback loops, and integration of extracellular cues, such as signaling pathways. Particularly, transcriptional switches during cell fate specification often involve specialized enhancers that control the ‘upstream TFs’, such as during neural tube patterning^83^. Such enhancers, that integrate multiple patterning cues, may be missed by the current enhancer modeling strategies. Indeed, CREsted and other CNN modeling frameworks are trained on enhancer logic across a set of similar/co-regulated enhancers, for example all mesenchymal enhancers downstream of TWIST1-ALX4 share the coordinator motif. Nevertheless, we believe that a comprehensive understanding of the ‘enhancer battery’ for each cell state, using TF-MINDI, is a first necessary step towards the inference of dynamic GRNs including repression and feedback loops.

In conclusion, we extracted human-interpretable *cis*-regulatory rules underlying early human neural development. These rules consist of cell type-specific TF binding motifs, their instances, affinity and combinations across cell type-specific chromatin accessibility peaks. We show that these rules are robust across biological systems, conserved across species and sufficient to generate novel enhancers with cell type-specific activity.

## Acknowledgments

This work was funded by the following grants to S. Aerts: ERCAdG (101054387); KU Leuven grant (C14/22/125), CZI (DI2-0000000068), Interuniversity BOF programme (IBOF/25/010) and Foundation Against Cancer (2020-1396 & 2024-140) and the following grants to S. Linnarson: Erling-Persson Foundation Human Developmental Cell Atlas grant, Knut and Alice Wallenberg Foundation grants 2015.0041, 2018.0172 and 2018.0220, Swedish Foundation for Strategic Research SB16-0065, and EU Horizon2020 BRAINTIME project 874606 and PhD fellowship from FWO to S. De Winter (1191323N).

We thank Lukas Mahieu for the co-development of the TF-MINDI Python package. We thank Darina Abaffyová for proofreading the manuscript and providing comments and for discussions about implementation details of TF-MINDI. We thank Peter Lönnerberg for preprocessing the human embryo data and making it available through EGA. We also thank Prof. Dr. Tatjana Sauka-Spengler and Dr. Ruth Williams for their support with helping to set up the chicken embryo enhancer reporter assays. Finally, we thank Professor Adrian Ranga, Dr. Abdel Rahman Abdel Fattah and Dr. Brian Daza for their help with setting up the neural tube organoid protocol. Computing was performed at the Vlaamse Supercomputer Center and VIB datacore.

## Author contribution statement

● Conceptualization: S.D.W & S.A.
● Experiments and sample preparation: S.D.W., V.C., L.H., C.M., & R.V.P.
● Computational analysis: S.D.W. & C.M.
● Writing of original draft: S.D.W., C.M. & S.A.
● Review & editing: S.D.W., C.M. & S.A.

## Competing interest statement

There are no competing interests to declare.

## Methods

### Experimental model and study participant details

#### Collection of human fetal samples

The experiments on human embryos in this study fall under category 2 as defined by the International Society for Stem Cell Research (ISSCR) 2021 guidelines. All experiments followed relevant guidelines and regulations, including the ISSCR 2021 guidelines. Human fetal samples were collected from routine termination of pregnancies at the Karolinska University Hospital following informed consent of the donors. The donors were informed of the purpose of the research and specific procedures the tissues will be used for. The use of fetal samples collected from abortions was approved by the Swedish Ethical Review Authority and the National Board of Health and Welfare (Etikprövningsmyndigheten; DNR2020-02074). Embryos were not kept in culture and were immediately dissected after collection. The embryo was staged according to the Carnegie stages. Following the dissection, the samples were transferred to ice-cold Hibernate E medium (ThermoFisher, A1247601) and processed the same day.

#### Generation of human neural tube organoids

The generation of human neural tube organoids falls under category 1a as defined by the ISSCR 2021 guidelines. Two induced Pluripotent Stem Cell lines (iPSC): IPSC EPITHELIAL-1 (RRID:CVCL_EE38), Purchased from Sigma-Aldrich (iPSC0028), and hereafter referred to as Sigma iPSC, and BJ1 iPSC (iPSC generated from BJ fibroblasts (RRID:CVCL_3653) and kindly gifted by professor Catherine Verfaillie (Leuven Stem Cell Institute) were used along with one human Embryonic Stem Cell (ESC) line (H9-ESC; RRID:CVCL_9773), kindly gifted by professor Pierre Vanderhaeghen (VIB Center for Brain and Disease research). The identities of the cell lines were validated based on their genomic sequences (see methods details below). Pluripotent stem cells were cultured in flat bottom, TC-treated 6 well plates (Avantor; 734-2323). For culture details see methods section below. All human iPSC and ESC experiments were approved by the Ethical Committee of KU Leuven (ECD S63316). Human neural tube organoids were generated based on Fattah *et al.* 2021 Nature Communications^39^. At 60-70% confluency pluripotent stem cells were washed twice using Phosphate Buffered Saline (PBS; ThermoFisher, 14190169; 2ml) next a single cell suspension was made by adding 1ml of room temperature TrypLE Express Enzyme (ThermoFisher, 12605010) and incubation for 3 minutes at 37°C followed by pipetting up and down using a 1ml pipette. Immediately after the TrypLE reaction was inhibited by adding 2 ml of DMEM/F12 FBS (Fetal Bovine Serum (FBS; ThermoFisher, A5256701) 20% V/V; Penicillin-Streptomycin (ThermoFisher, 15140122) 50µg/ml in DMEM/F12 (ThermoFisher, 11320033)). Cells were pelleted at 300 RCF for 3 minutes and, after counting on a LUNA-FL Dual Fluorescence Cell Counter resuspended, to 3.5e6 cells per ml in GM_0_ (basal GM (N-2 Supplement (ThermoFisher, 17502048) 1x, B-27 Supplement (ThermoFisher, 17504044) 1x, GlutaMax Supplement (ThermoFisher, 35050061) 1x, Sodium Pyruvate (ThermoFisher, 11360070) 1mM, MEM Non-Essential Amino Acids Solution (ThermoFisher, 11140050) 1x, Penicillin-Streptomycin (ThermoFisher, 15140122) 50 µg/ml, Neurobasal Medium (ThermoFisher, 21103049) 50% V/V and DMEM/F12 (ThermoFisher, 11320033) 50% V/V) with RHO/ROCK inhibitor (Y-27632 (Dihydrochloride), Stemcell technologies, 100-1044) 1µM). Next, the polyethylene glycol A (PEG-A) hydrogel matrix (see Methods detail below) was prepared by first adding Corning Mouse Laminin (Avantor, 734-1098) 0.1 mg per ml followed by activated factor 13 (see Methods details below) in that order to a PEG mixture (see Methods details below; PEG 1.5 % or 2 % V/V, buffer (TRIS 0.05 mM, 𝐶𝑎𝐶𝑙2 0.05 mM; pH 7.6; filter sterilized) in 𝐺 𝑀0 medium). After mixing the matrix by pipetting up and down, without generating bubbles, for 1-2 minutes 3.5e4 cells per µl mixture were added. The matrix and gell mixture was mixed gently one more time by pipetting and 10 µl droplets were generated in each well of a TC-treated 96 well plate (Corning, 3603). When the matrix started to gell the plate was turned up-side down for 10 seconds, after which it was turned right-side up for 10 seconds. This was repeated for a total of 5 times, after which the plates were kept at room temperature. After 20 minutes of gelling 200 µl of GM_0_ medium was added to each well and the organoids were incubated at 37 °C 5 % CO_2_ for a total of 11 days, or until the timepoint of single cell multiome. At day 3 of differentiation medium was changed with basal GM supplemented with retinoic acid (Sigma-Aldrich, R2625-100MG) 0.25 µM and Smoothened agonist (Sigma-Aldrich, 566661-500UG) 1 µM. At day 5, 7 and 9 the medium was changed with basal GM.

#### *In vitro* differentiation of facial mesenchyme/cranial neural crest cells

In vitro differentiation of facial mesenchyme/cranial neural crest cells falls under category 1a as defined by the ISSCR 2021 guidelines. Cells were differentiated as previously described^84–86^ At 60-70% confluency sigma iPSCs were washed twice using Phosphate Buffered Saline (PBS; ThermoFisher, 14190169) and detached from the culture plate using Collagenase (ThermoFisher, 17104019) treatment at 37°C for 45min-1.5h, until small colonies have released and large colonies have rolled up edges. Colonies of cells were collected by careful aspiration of Collagenase and adding 4ml PBS to each well to transfer the colonies to a 15ml tube followed by washing and scraping (using pipette tip) the wells a final time to come to a total of 5-8ml PBS with Colonies. Finally, colonies were allowed to settle to the bottom of the tube (1-2min) and PBS was aspirated carefully. Colonies were disaggregated to cell aggregates of around 100 cells in size by adding 1ml of NCC differentiation medium (bFGF (ThermoFisher, 100-18B-50UG) 20 ng/ml, Insulin (Sigma-Aldrich, 11070-73-8) 5e3 ng/ml, EGF (ThermoFisher, AF-100-15-500UG) 20ng/ml in Neural Crest Base Medium (B27 (ThermoFisher, 17504044) 0.5x, N2 (ThermoFisher, 17502048) 0.5x, Glutamax (ThermoFisher, 35050061) 1x, Penicillin-Streptomycin (ThermoFisher, 15140122) 50 µg/ml, DMEM F12 (ThermoFisher, 11320033) 1x, Neurobasal medium (ThermoFisher, 21103049) 1x)) and pipetting once using a P1000 pipette, waiting for a small pellet to form and pipetting that pellet once more. Next, another 1ml of NCC differentiation medium was added and cell aggregates were transferred to a 10cm dish, distributed evenly and left O/N at 37°C. For the first three following days, the medium was changed as needed to remove dead cells by gently pipetting the cell aggregates using a 5ml pipette to a 15ml tube and waiting for them to settle to the bottom after which the supernatant is removed and cell aggregates are resuspended in 2ml NCC differentiation medium and transferred to a 10 cm plate containing 1ml pre-warmed NCC differentiation medium. At day four, the medium is changed in a similar fashion and the cells are transferred to a TC treated tissue culture plate where they are kept for at least 3 more days until the cell aggregates start to attach to the culture plate. If cells are not attached after 3 days the medium is changed as before, without changing plates. Once the cells are attached the medium is changed once more. Three days later, when cells cover the entire tissue culture plate, cells are transferred to a fibronectin coated plate. For this, tissue culture plates are coated with Fibronectin (Sigma-Aldrich, FC010) 7.5µg/ml in PBS for at least 30min at room temperature. Next, cells are washed once using PBS and detached using Accutase (ThermoFisher, 00-4555-56) 60% (V/V) in PBS for 5 min at 37°C. Accutase reaction is inhibited using double the volume of NCC Maintenance Medium (bFGF (ThermoFisher, 100-18B-50UG) 20 ng/ml, EGF (ThermoFisher, AF-100-15-500UG) 20ng/ml, BSA 1mg/ml in Neural Crest Base Medium). Next, cell aggregates are transferred to fibronectin coated plate in pre-warmed NCC maintenance medium and allowed to attach at °C for 15-30min after which medium is replaced with NCC maintenance medium. To passage the cells, medium is removed without tilting the plate, cells are washed once using PBS and are detached using Accutase 60% (V/V) tilting the plate to distribute the solution and incubation for a total of 2-5min at 37°C until cells are detached (single cells). At 2 min, the plate is tilted again in all directions and returned to 37°C for 2-3min after which it is tilted again. Next, single cells are transferred to a fibronectin coated plate in pre-warmed NCC maintenance medium. From passage three onwards the same procedure is repeated but using Long Term Maintenance Medium instead (ChiRON 99021 (Sigma-Aldrich, SML1046-5MG) 3 µM, BMP2 (ThermoFisher, 120-02-10UG) 100 pg/ml in NCC Maintenance Medium).

#### Chicken embryos

Fertilized chicken eggs, obtained from ”Kwekerij van het Hallerbos” were incubated at 37 °C at 70% humidity for 30 hours to reach Hamburger-Hamilton (HH) stage 4. Afterwards, embryos were removed from the egg and kept in culture for an additional 24 hours.

### Method detail

#### Human neural tube organoid culture

##### Factor 13 activation

1U of reconstituted Thrombin (Sigma-Aldrich, 10602400001) in Ehrbars buffer (TRIS 10mM; NaCl 150mM; CaCl_2_ 25mM pH 7.4) per 90U of reconstituted Fibrogammin (UZ Leuven pharmacy) was mixed and put in water bath at 37 °C for 30 minutes while swirling every 5 minutes. Activated factor 13 was filter sterilized, aliquoted and stored at -80 °C.

##### PEG-A hydrogel precursor synthesis

PEG-A hydrogel precursors, as described in Ranga *et al*. nature communications^48^, were kindly gifted by professor Adrian Ranga (Department of Mechanical Engineering at KU Leuven).

##### Human embryonic/pluripotent stem cell culture

TC-treated well plates were coated using Geltrex Matrix (ThermoFisher, A1413301) as follows. Geltrex Matrix was thawed in ice overnight at 4 °C and diluted in DMEM/F12 (ThermoFisher, 11320033) 1 % V/V. A sufficient amount of diluted Geltrex Matrix was added to each well to coat the surface and the culture plates were incubated at 37 °C for a minimum of 60 minutes. The remaining Geltrex mixture was removed prior to culture. Human Pluripotent stem cells were cultured on the Geltrex Matrix coated well plates at 37 °C 5 % CO_2_ in Essential 8 Flex Medium (ThermoFisher, A2858501) supplemented with Penicillin Streptomycin (ThermoFisher, 15140122) 50 µg per ml. The medium was changed every other day. Cells were passaged when they reached 70 % by washing them twice using DPBS without Calcium and Magnesium (ThermoFisher, 14190) and adding EDTA (ThermoFisher, 15575020) 0.5 mM followed by incubation at 37 °C. After 3-4 minutes the EDTA solution was aspirated and replaced by Essential 8 Flex Medium. Cells were scraped using a cell scraper (Corning, 734-0385) and split in a 1:6 ratio.

##### Single cell suspension

PEG hydrogel droplets were removed from the 96 well plate by gently scraping with a 100 µl pipette tip and transferred to a tube on ice using a p1000 pipette by putting the pipette tip on the droplet and creating a gentle negative pressure, making sure that the droplet does not enter the pipette tip. Organoids were dissociated by adding room temperature TrypLE Express Enzyme (ThermoFisher, 12605010) and incubation at 37 °C while pipetting up and down until no large pieces of hydrogel were visible anymore or a maximum of 7.5 minutes. Immediately after the TrypLE reaction was inhibited by adding twice the volume of DMEM/F12 FBS (Fetal Bovine Serum (FBS; ThermoFisher, A5256701) 20 % V/V; Penicillin-Streptomycin (ThermoFisher, 15140122) 50 µg in DMEM/F12 (ThermoFisher, 11320033)). The single-cell suspension was pelleted at 500 RCF for 4 minutes, resuspended in DMEM/F12 FBS and counted on LUNA-FL Dual Fluorescence Cell Counter. Immediately after the cells were processed for 10x single-cell multiome.

#### Human neural tube organoid multiome

##### 10x multiome nuclei isolation

To isolate nuclei from the single cells of the dissociated neural tube organoids, protocol CG000365 (10x Genomics) was followed. For each developmental time point, 500k cells were resuspended in 100 μl nuclei lysis buffer (10mM Tris-HCl pH 7.4; 10mM NaCl; 3mM MgCl_2_; 0.1 % Tween-20; 0.1 %NP40; 0.01 % Digitonin; 1 % BSA; 1mM DTT and 1U per µl RNasin ribonuclease inhibitor in nuclease-free water). After 5 min incubation on ice, 1ml chilled wash buffer was added to the lysed cells (10mM Tris-HCl pH 7.4; 10mM NaCl; 3mM MgCl_2_; 0.1 % Tween-20; 1 % BSA; 1mM DTT and 1U per µl RNasin ribonuclease inhibitor in nuclease-free water). Nuclei were pelleted by centrifugation at 500 g for 5 min at 4 °C and resuspended in a 1x nuclei buffer (10x Genomics) supplemented with 1mM dithiothreitol and 1 U/µl RNasin ribonuclease inhibitor. Nuclei suspensions were passed through a 40 µm Flowmi filter (VWR Bel-Art SP Scienceware). Nuclei concentration was assessed using the LUNA-FL Dual Fluorescence Cell Counter.

##### 10x multiome library preparation

10x multiome library preparation Single-cell libraries were generated using the 10X Chromium Single-Cell Instrument and NextGEM Single Cell Multiome ATAC+Gene Expression kit (10X Genomics) according to the manufacturer’s protocol. In brief, single nuclei of neural tube organoids were incubated for 60 min at 37 °C with a transposase that fragments the DNA in open regions of the chromatin and adds adapter sequences to the ends of the DNA fragments. After generation of nanoliter-scale gel bead-in-emulsions (GEMs), GEMs were incubated in a C1000 Touch Thermal Cycler (Bio-Rad) under the following program: 37 °C for 45 min, 25 °C for 30 min, and hold at 4 °C. Incubation of the GEMs produced 10x barcoded DNA from the transposed DNA (for ATAC) and 10x barcoded, full-length cDNA from poly-adenylated mRNA (for GEX). This was followed by a quenching step that stopped the reaction. After quenching, single-cell droplets were broken and the transposed DNA and full-length cDNA were isolated using Cleanup Mix containing Silane Dynabeads. To fill gaps and generate sufficient mass for library construction, the transposed DNA and cDNA were amplified via PCR: 72 °C for 5 min; 89 °C for 3 min; seven cycles of 98 °C for 20s, 63 °C for 30 s, 72 °C for 1 min; 72 °C for 1 min; and hold at 4 °C. The pre-amplified product was used as input for both ATAC library construction and cDNA amplification for gene expression library construction. Illumina P7 sequence and a sample index were added to the single-strand DNA during ATAC library construction via PCR: 98 °C for 45 s; 7–9 cycles of 9 °C for 20 s, 67 °C for 30 s, 72 °C for 20 s; 72 °C for 1 min; and hold at 4 °C. The sequencing-ready ATAC library was cleaned up with SPRIselect beads (Beckman Coulter). Barcoded, full-length pre-amplified cDNA was further amplified via PCR: 98 °C for 3 min; 6–9 cycles of 98 °C for 15 s, 63 °C for 20 s, 72 °C for 1 min; 72 °C for 1 min; and hold at 4 °C Subsequently, the amplified cDNA was fragmented, end-repaired, A-tailed, and index adaptor ligated, with SPRIselect cleanup in between steps. The final gene expression library was amplified by PCR: 98 °C for 45 s; 5–16 cycles of 98 °C for 20s, 54 °C for 30 s, 72 °C for 20 s, 72 °C for 1 min; and hold at 4 °C. The sequencing-ready GEX library was cleaned up with SPRIselect beads.

##### 10x multiome sequencing

Before sequencing, the fragment size of every library was analyzed using the Bioanalyzer high-sensitivity chip. All 10x single cell Multiome ATAC libraries were sequenced on NextSeq2000 instrument (Illumina) or on NovaSeq6000 instrument (Illumina) with the following sequencing parameters: 51 bp read 1, 8 bp index 1, 24 bp index 2, 51 bp read 2. All 10x single cell Multiome Gene Expression libraries were sequenced on NextSeq2000 instrument (Illumina) or on NovaSeq6000 instrument (Illumina) with the following sequencing parameters: 28 bp read 1, 10 bp index 1, 10 bp index 2, 90 bp read 2.

#### Human embryo sample preparation

##### 10x multiome library preparation

Tissue was gently minced using a razor blade and cells were dissociated using a papain dissociation kit (Worthington) with 200 U per ml DNAse at 37 °C for 10 min following the manufacturer’s provided protocol. The suspension was triturated using a glass pipette to dissolve remaining clumps before being filtered through a 30 µm filter (CellTrics). After dissociation, the cells were washed with 1X EBSS and concentrated (200g, 5 min). 1e6 cells were pelleted (500g, 5 min) in a 2 ml LoBind Eppendorf tube. Nuclei were then isolated from the pellet by incubating them with 100 µl dissociation mix (0.001 % digitonin, 0.01 % NP40, 1 mM dithiothreitol, 1 U per µl RNAse inhibitor, 0.1 % Tween-20, 1 % BSA, 10 mM Tris-HCl, 10 mM NaCl, 3 mM MgCl_2_) for 5 min before adding 1 ml wash buffer (1 mM dithiothreitol, 1 U per µl RNAse inhibitor, 0.1 % Tween-20, 1 % BSA, 10 mM Tris-HCl, 10 mM NaCl, 3 mM MgCl_2_), pelleting the isolated nuclei (500g, 5 min) and resuspending in 1X nuclei buffer (10X Genomics). Single-cell libraries were generated using the 10X Chromium Single-Cell Instrument and NextGEM Single Cell Multiome ATAC+Gene Expression kit (10X Genomics) according to the manufacturer’s protocol as described in the human neural tube organoids section. In total, 4 ATAC and 4 Gene Expression libraries were generated for the embryonal sample.

##### 10x multiome library sequencing

The fragment size of the libraries were analyzed using the Bioanalyzer high-sensitivity chip. All embryonal libraries were first sequenced on the NextSeq2000 instrument (Illumina) before being sequenced deeper on the NovaSeq6000 instrument (Illumina). The ATAC libraries were sequenced with the following parameters: 50 bp read 1, 8 bp index 1, 24 bp index 2, 49 bp read 2. The Gene Expression libraries were sequenced with the following parameters: 28 bp read 1, 10 bp index 1, 10 bp index 2, 90 bp read 2.

#### Human neural tube data analysis

##### Samples

Eight individual single-cell multiome libraries were generated:

● A single library containing a mix of organoids at 1, 3, 5, 7, 9 and 11 days of differentiation cultured in 1.5 % PEG hydrogel matrix: IPS 081787 575955 10x_multiome_neural_tube_organoids_1_5_PEG.
● A single library containing a mix of organoids at 1, 3, 5, 7, 9 and 11 days of differentiation cultured in 2 % PEG hydrogel matrix: IPS c13535 a58829 10x_multiome_neural_tube_organoids_2_PEG.
● Two libraries containing a mix of organoids at 1, 2, 3, 6 and 7 days of differentiation cultured in 2 % PEG hydrogel matrix:6-7_1-2-3_1 and 6-7_1-2-3_2.
● A single library containing a mix of organoids at 6 and 7 days of differentiation cultured in 2 % PEG hydrogel matrix: 6-7.
● A single library containing a mix of organoids at 4, 5, 8 and 9 days of differentiation cultured in 2 % PEG hydrogel matrix: 8-9_4-5.
● A single library containing a mix of organoids at 8 and 9 days of differentiation cultured in 2 % PEG hydrogel matrix: 8-9.
● A single library containing a mix of organoids at 10 and 11 days of differentiation cultured in 2 % PEG hydrogel matrix:10-11.

For each sample a mix of H9-ESC, Sigma iPSC and BJ1 iPSC cells were used.

##### RNA count matrices and ATAC fragment files

single-cell RNA count matrices and single-cell ATAC fragment files were generated for each sample using the Cell Ranger ARC (v2.0.2;RRID:SCR_02389) command cellranger-arc count command using as the GRCh38 reference genome supplied with Cell Ranger ARC (v2020-A-2.0.0).

##### Cell line demultiplexing

Variants were called from in house Omni-ATAC-seq^87^ data that was performed on each cell line individually (unpublished data). First, bam files per cell line were merged using samtools^88^ (v1.11; RRID:SCR_002105) merge, sorted using samtools sort and indexed using samtools index. Next, for each cell line merged bam file common variants were called using the bcftools^89^ (v1.11; RRID:SCR_005227) mpileup command using common short genetic variants from 1000 Genomes project as targets parameter (-T) and 8,000 as max depth (-D) parameter followed by the call command from bcftools. Next, single-cell multiome samples were demultixpled per cell line using popscle (RRID:SCR_026707) demuxlet^90^ after first sorting the variant VCF file as the single-cell multiome sample bam file, using a custom script *sort_VCF_same_as_bam*, filtering the sample bam file to only retain reads that overlap with SNPs in the VCF file and have a valid barcode passing the Cell Ranger ARC quality control, using a custom script *filter_bam_file_for_popscle_dsc_pileup*, (custom scripts can be found in the popscle helper tools GitHub repository: https://github.com/aertslab/popscle helper tools) and creating a pileup of reads for each SNP in the VCF file and each cell barcode using the single-cell multiome sample bam file and finally popscle demuxlet was run on the sample pileup, variant VCF file using GT as field (–field) parameter.

##### Single-cell RNA quality control

Cells with less than 3 genes expressed and genes expressed in less than 200 cells were removed using the Scanpy^91^ (v1.9.1; RRID:SCR_018139) functions filter cells and filter genes. Next, quality control metrics were calculated using the Scanpy function calculate qc metrics using qc vars = [”mt”], percent top = None and log1p = False as parameters. Cells with less than 25 % mitochondrial counts, more than a total of 2,000 counts and less than a total of 9,000 genes with counts were kept resulting in a total of 27,552 cells passing quality control.

##### Single-cell RNA batch effect correction

Batch effect correction was performed using harmonypy^92^ (v0.0.1; RRID:SCR_022798) using as batch key the combination of experimental batch and cell line on 50 PCA coordinates calculated using the Scanpy^91^ (v1.9.1; RRID:SCR_018139) function pca.

##### Single-cell RNA cell type annotation

Cells were first annotated as Facial mesenchyme, Neural crest, Neuron, Peripheral neuron, Pluripotent stem cell or neural progenitor by first normalizing, log transforming and scaling the gene expression matrix using the Scanpy^91^ (v1.9.1; RRID:SCR_018139) functions *normalize_per_cell*, *log1p* and *scale* and clustering the cells using the Leiden algorithm^93^ at a resolution of 5.1 (making use of the leiden function from Scanpy), calculating the median expression per cluster, binarizing this matrix using a gene expression threshold of 0, and calculating the jaccard distance using Scipy^94^ (v1.9.3; RRID:SCR_008058) between the binarized median expression matrix and a knowledge matrix where containing marker genes per cell type and where the value is 1 for markers of that cell type and 0 otherwise constructed from the marker gene table (**Table S1**). For each cluster, the annotation with minimal distance was selected, breaking ties by selecting a random annotation with equal distance. neural progenitors were further annotated as neural progenitor, neural progenitor early neuron, KRT19+ neural progenitors, FOXG1+ neural progenitors, NFE2L3+ neural progenitors and KRT19+/NFE2L3+ neural progenitors using the same procedure at a Leiden resolution of 1.9 after subsetting for neural progenitors and re-normalizing, scaling and integrating (see batch effect correction) the data. Neural crest cells were further annotated as pre-migratory neural crest, bonafide neural crest, EMT neural crest, mesenchyme and peripheral neuron manually based on the expression of marker genes. Neuron cells and neural progenitor early neuron cells were further annotated as ASCL1+ Neurons, SST+ Neurons, SOX2+ Neurons, ASCL1+ Neurons, UNCX+ Neurons, GAD1+ Neurons and GATA2+ Neurons based on the expression of marker genes.

##### Single-cell ATAC consensus peaks

A set of consensus peaks was called by first generating pseudobulk ATAC-seq profiles for each of the highest resolution cell types (see single-cell RNA cell type annotation) using the export pseudobulk function, with default parameters, from pycisTopic^52^ (v1.0.3.dev20+g8955c76, RRID:SCR_026618) followed by peak calling using MACS^95^ (v2.2.9.1; RRID:SCR_013291) using the pycisTopic function peak calling, using default parameters, and consensus peak generation using the pycisTopic function get consensus peaks, using a peak width of 500 bp and default parameters. This resulted in 488,211 consensus peaks.

##### Single-cell ATAC quality control

Sample and barcode level quality control data was calculated using the *compute_qc_stats* function from pycisTopic^52^ (v1.0.3.dev20+g8955c76, RRID:SCR_026618) cells were filtered based on the number of unique fragments per cell, Fraction of Reads in Peaks (FRIP) and transcription start site (TSS) enrichment.

Following thresholds were used for each sample:

● sample IPS 081787 575955 10x_multiome_neural_tube_organoids_1_5_PEG
  ○ log unique number of fragments: > 3.5
  ○ FRIP: > 0.5
  ○ TSS enrichment: > 5
● sample IPS c13535 a58829 10x_multiome_neural_tube_organoids_2_PEG
  ○ log unique number of fragments: > 3.5
  ○ FRIP: > 0.5
  ○ TSS enrichment: > 5
● sample 10-11
  ○ log unique number of fragments: > 3.25
  ○ FRIP: > 0.45
  ○ TSS enrichment: > 6
● sample 6-7
  ○ log unique number of fragments: > 3.5
  ○ FRIP: > 0.45
  ○ TSS enrichment: > 6
● sample 6-7_1-2-3_1
  ○ log unique number of fragments: > 3.75
  ○ FRIP: > 0.5
  ○ TSS enrichment: > 7
● sample 6-7_1-2-3_2
  ○ log unique number of fragments: >3.75
  ○ FRIP: > 0.5
  ○ TSS enrichment: > 6
● sample 8-9
  ○ log unique number of fragments: > 3.25
  ○ FRIP: > 0.4
  ○ TSS enrichment: > 7
● sample 8-9_4-5
  ○ log unique number of fragments: > 3.75
  ○ FRIP: > 0.5
  ○ TSS enrichment: > 6

Cells that were predicted as doublets by demuxlet (see: Cell line demultiplexing) were also filtered out resulting in a total of 28,705 cells passing quality control.

##### Single-cell ATAC topic modeling

Four topic models were generated: one on all cells, one on the subset of cells annotated as neural progenitors, one on the subset of cells annotated as neuron (Neuron cells and neural progenitor early neuron cells) and one on the subset of cells annotated as neural crest. Topic modeling was performed using Mallet^96^ (v202108; RRID:SCR_026708) using the pycisTopic^52^ (v1.0.3.dev20+g8955c76, RRID:SCR_026618) function *run_cgs_models* mallet using 500 iterations, alpha equal to 50, eta equal to 0.1, alpha by topic equal to True and eta by topic equal to False and using random seed 555. Topic modeling was performed separately for all cells and for each subset of cells. Model evaluation was performed using the pycisTopic function evaluate model that calculates model coherence^97^, log-likelihood^98^, model density^99^ and a divergence-based metric^100^. A model of 50 topics was selected for the model containing all cells, a model of 30 topics was selected for the model containing neural progenitors, a model of 25 topics was selected for the model containing neurons and a model of 20 topics for the model containing neural crest cells. The region topic probability matrix was thresholded using the *binarize_topics* function from pycisTopic using the otsu method^101^ and the cell topic probability matrix using the same function using the li method^102^.

##### Sequence-to-function model training: DeepNeuralTube°

A deepTopic model^19^, now implemented in CREsted^24^ (RRID:SCR_026617), containing 5 convolutional and max pooling layers: dimensions (500, 1,024), (125, 512), (32, 512), (8, 512) and (2, 512) a fully connected dense layer of dimension 1,024 and a dense output layer of dimension 75 was trained to classify consensus peaks in topics based on their genomic sequence. GELU was used as activation function, 17 as kernel size, 4 as max pooling size, a dropout of 0.15 was used for the convolutional layers, a dropout of 0.5 was used for the dense layers, the convolutional layers were l2 normalized using an epsilon value of 1e-6 and the dense layers were l2 normalized using an epsilon value of 1e-3. The Adam optimizer was used with a learning rate of 0.001 that was dynamically scaled and binary cross entropy as a loss function and a batch size of 1,024. The model was built and trained using Tensorflow (v2.4.1; RRID:SCR_016345). As training data all binarized topics (see single-cell ATAC topic modeling) from the subset of neural progenitors, neurons and neural crest together with 10 topics form the topic model containing all cells representing the pluripotent stem cell state were used for a total of 75 classes/topics. The data was augmented by training on both forward and reverse complement sequence and 10 % of the data was used for validation and 10 % for testing for a total of 5,192,175 training samples, 650,280 test samples and 647,805 validation samples. Based on the validation area under the precision recall curve and validation area under the receiver operator curve the model after 23 epochs was selected.

The classes of the model correspond as follows to the different data subsets:

● 0 ≤ class_index_ < 25: neuron
● 25 ≤ class_index_ < 55: neural progenitor
● 55 ≤ class_index_ < 65: Neural crest
● 65 ≤ class_index_ < 75: pluripotent stem cell

#### Human embryo data analysis

##### RNA count matrices and ATAC fragment files

single-cell RNA count matrices and single-cell ATAC fragment files were generated using the Cell Ranger ARC (v2.0.0; RRID:SCR_023897) command cellranger-arc count using GRCh38.p13 Gencode V35 primary sequence assembly as a reference genome as described in Braun *et al*.^51^.

##### Single-cell RNA quality control

Cells with less than 3 genes expressed and genes expressed in less than 200 cells were removed using the Scanpy^91^ (v1.8.2; RRID:SCR_018139) functions *filter_cells* and *filter_genes*. Next, quality control metrics were calculated using the Scanpy function *calculate_qc_metrics* using qc vars=[”mt”], percent top = [20] and log1p = True as parameters. Cells were filtered if they were more than 5 median absolute distances (MADs) away from the median log1p transcript count, log1p number of genes or percentage counts in top 20 genes, or more than 3 MADs away from the median mitochondrial fraction, resulting in 34,453 high quality cells. Doublets were detected using Scrublet^103^ (RRID:SCR_018098) with a threshold of 0.2 and expected doublet rate of 6%. Counts were normalized to 10,000, log transformed and scaled. The top 20,000 highly variable genes were identified using the highly variable genes function using flavor=’seurat’. The top 50 PCA components were used to calculate KNN neighbors and leiden clustering at a resolution of 0.3.

##### Single-cell RNA cell type annotation

Clusters that were TUBB3+, ELAVL3+, INA+, STMN2+ and MAP2+ were annotated as neurons, with the additional expression of *SIX1* indicating peripheral neurons. *SOX2*, *PAX6*, *RFX4* and *PTPRZ* were used to annotate neural progenitor cells. The neural crest was identified based on the expression of *SOX10*, *TFAP2A*, *TFAP2B*, *PAX3* and *GRHL2*. *FOXC1*, *FOXC2* and *TWIST1* were used as mesenchymal markers and *FLT1*, *PTPRM* and *CLDN5* conversely were used to annotate endothelial cells. *OTOGL* and *OC90* marked the Otic placode and microglia were based on *RUNX1* and *CSF1R.* Neurons and neural progenitors were subclustered and neuronal subtypes were annotated based on the expression of *ASCL1*, *UNCX*, *GAD1*, *GATA2*, *SST* and *SOX2*. Conversely, the floor and roof plate were identified among the neural progenitor cells based on the expression of *SHH* and *MSX1* respectively. Conversely, regionalization was assessed based on expression of *FOXG1/EMX2* (forebrain), *OTX2/EN1* (midbrain), *PAX8* (hindbrain) and *HOXB3* (spinal cord).

##### Single-cell ATAC consensus peaks

The consensus peak set derived from neural tube organoids was used.

##### Single-cell ATAC quality control

Sample and barcode level quality control data was calculated using the pycisTopic^52^ (v2.0a, RRID:SCR_026618) command pycistopic qc, using a TSS annotation bed file that was generated using the pycisTopic command pycistopic tss get_tss using hsapiens_gene_ensembl as biomart name. High quality cells were selected based on the number of unique fragments per cell and TSS enrichment. The minimal number of unique fragments per cell was automatically determined using otsu thresholding^101^ and a TSS enrichment threshold of 7 was used. This resulted in 38,998 high quality cells passing quality control.

##### Single-cell ATAC topic modeling

Three topic models were generated: one on the subset of cells annotated as neural progenitors, one on the subset of cells annotated as neuron and one on the subset of cells annotated as neural crest. First, a fragment count matrix was generated for each sample using the pycisTopic^52^ (v2.0a, RRID:SCR_026618) function create_fragments_matrix_from fragments, this matrix was binarized (>0), merged in one count matrix using the Scipy^94^ (v1.15.2; RRID:SCR_008058) function sparse.hstack and subsequently split in a matrix for each of neural progenitors, neurons and neural crest cells and saved as matrix market format. A mallet corpus file was generated for each binary matrix using the pycisTopic command pycistopic topic_modeling mallet create_corpus and finally topic modeling was run using Mallet^96^ (v202108; RRID:SCR_026708) using the pycisTopic command: pycistopic topic_modeling mallet run using 300 iterations and default parameters otherwise. Model evaluation was performed using the pycisTopic command pycistopic topic_modeling mallet stats that calculates model coherence^97^, log-likelihood^98^, model density^99^ and a divergence-based metric^100^. A model of 60 topics was selected for the model containing neural progenitors, a model of 30 topics was selected for the model containing neurons and a model of 30 topics for the model containing neural crest cells. The region topic probability matrix was thresholded using the binarize topics function from pycisTopic using the otsu method^101^ and the cell topic probability matrix using the same function using the li method^102^.

##### Sequence-to-function model training: DeepNeuralTube^e^

A deepTopic model^19^, now implemented in CREsted^24^ (RRID:SCR_026617), containing 5 convolutional and max pooling layers: dimensions (500, 1,024), (125, 512), (32, 512), (8, 512) and (2, 512) a fully connected dense layer of dimension 1,024 and a dense output layer of dimension 120 was trained to classify consensus peaks in topics based on their genomic sequence. GELU was used as activation function, 17 as kernel size, 4 as max pooling size, a dropout of 0.15 was used for the convolutional layers, a dropout of 0.5 was used for the dense layers, the convolutional layers were l2 normalized using an epsilon value of 1e-6 and the dense layers were l2 normalized using an epsilon value of 1e-3. The Adam optimizer was used with a learning rate of 0.001 that was dynamically scaled and binary cross entropy as a loss function and a batch size of 1,024. The model was built and trained using Tensorflow (v2.4.1; RRID:SCR_016345). As training data all binarized topics (see single-cell ATAC topic modeling) from the subset of neural progenitors, neurons and neural crest together were used for a total of 120 classes/topics. Data was augmented by training on both forward and reverse complement sequence and 10 % of the data was used for validation and 10 % for testing for a total of 11,935,075 training samples, 1,494,835 test samples and 1,497,350 validation samples. Based on the validation area under the precision recall curve and validation area under the receiver operator curve the model after 36 epochs was selected.

The classes of the model correspond as follows to the different data subsets:

● 0 ≤ class_index_ < 30: neuron
● 30 ≤ class_index_ < 90: neural progenitor
● 90 ≤ class_index_ < 120: Neural crest

### Cell cycle annotation

Marker genes of the S-phase and G2 and M phase of the cell cycle^104^ were used in the Scanpy^91^ (v1.11.0; RRID:SCR_018139) function score genes cell cycle.

### Annotating accessibility topics to cell types

The Pearson correlation between the cell-topic-probabilities of all accessibility topics and the log CPM gene expression values of known marker genes using the pearsonr function from Scipy^94^ (v1.9.3; RRID:SCR_008058). Topics that had a correlation coefficient of at least 0.35 for any marker gene were kept for subsequent analyses.

### Validation of DeepNeuralTube sequence-to-function models based on Vista enhancers

Data was downloaded from the VISTA Enhancer browser^53,54^ (https://gitlab.com/egsb-mfgl/vistadata/-/raw/main/experiments.tsv.gz) and the DNA sequence of each element was obtained either from the human genome (hg38) or the mouse genome (mm10) dependent on the origin of each sequence. In case a sequence was shorter than 500 bp (the input size of DeepNeuralTube) the sequence was extended to 500 bp based on the genomic coordinates. In case the sequence was larger than 500 bp a sliding window, with step size of 1, was produced over the sequence. A prediction score for each sequence and using both DeepNeuralTube° and DeepNeuralTube^e^ was calculated and per experiment and output class of the model the maximum prediction score was retained. Ground truth neural tube enhancers were defined as enhancers that were positive in the tissue ”nt”, in at least 3 embryos and at least 50 % of tested embryos. Ground truth facial mesenchyme enhancers were defined as enhancers that were positive in the tissue ”fm”, in at least 3 embryos and at least 50 % of tested embryos. For both neural tube enhancers and facial mesenchyme enhancers precision-recall curves and receiver-operator curves were calculated using respectively the function metrics.precision_recall_curve and metrics.roc_curve from Scikit-learn^105^ (v1.5.2; RRID:SCR_002577) using the maximum prediction score across classes related to neural tube respectively facial mesenchyme in both organoid and embryo. We consider classes related to neural tube in the organoid as classes of the following topics: neural crest Topic 5, neural crest Topic 7, neuron Topic 1, neuron Topic 2, neuron Topic 3, neuron Topic 4, neuron Topic 6, neuron Topic 10, neuron Topic 11, neuron Topic 12, neuron Topic 13, neuron Topic 15, neuron Topic 16, neuron Topic 18, neuron Topic 19, neuron Topic 20, neuron Topic 21, neuron Topic 23, neuron Topic 24, neuron Topic 25, progenitor Topic 1, progenitor Topic 3, progenitor Topic 8, progenitor Topic 9, progenitor Topic 11, progenitor Topic 13, progenitor Topic 14, progenitor Topic 16, progenitor Topic 19, progenitor Topic 21, progenitor Topic 23, progenitor Topic 24, progenitor Topic 25, progenitor Topic 29 and progenitor Topic 30. We consider classes related to neural tube in the embryo as classes of the following topics: neuron Topic 1, neuron Topic 3, neuron Topic 5, neuron Topic 6, neuron Topic 7, neuron Topic 8, neuron Topic 9, neuron Topic 10, neuron Topic 11, neuron Topic 12, neuron Topic 13, neuron Topic 14, neuron Topic 15, neuron Topic 17, neuron Topic 18, neuron Topic 19, neuron Topic 22, neuron Topic 24, neuron Topic 26, neuron Topic 27, neuron Topic 29, neuron Topic 30, progenitor Topic 1, progenitor Topic 3, progenitor Topic 4, progenitor Topic 8, progenitor Topic 10, progenitor Topic 11, progenitor Topic 16, progenitor Topic 17, progenitor Topic 21, progenitor Topic 22, progenitor Topic 28, progenitor Topic 29, progenitor Topic 31, progenitor Topic 32, progenitor Topic 36, progenitor Topic 40, progenitor Topic 41, progenitor Topic 44, progenitor Topic 45, progenitor Topic 49, progenitor Topic 51, progenitor Topic 57, progenitor Topic 58, progenitor Topic 59, neural crest Topic 12, neural crest Topic 13, neural crest Topic 15. We consider classes related to facial mesenchyme in the organoid as classes of the following topic: neural crest Topic 3. We consider classes related to facial mesenchyme in the organoid as classes of the following topics: neural crest Topic 1, neural crest Topic 19, neural crest Topic 21 and neural crest Topic 29.

### Motif enrichment analysis

A cistarget motif ranking database was created by first getting the genomic sequence of each organoid consensus peak with 1,000 bp background padding using the create_fasta_with_paddedbg_from_bed script from the create_cisTarget_databases repository (https://github.com/aertslab/) and scoring each DNA sequence using Cluster-Buster^106^ (RRID:SCR_024810) using the create_cisTarget_databases command create_cistarget_motif_databases followed by the generation of the ranking databases using the command convert_motifs_or_tracks_vs_regions_or_genes_scores_to_rankings_ cistarget_dbs. Next, the SCENIC+^52^ (v1.0a2; RRID:SCR 026702) command scenicplus grn_inference motif_enrichment_cistarget command was used to perform cisTarget motif enrichment analysis for each organoid and embryo topic.

### *De novo* motif discovery using TF-MoDISCo

For all organoid and embryo topics using respectively DeepNeuralTube° and DeepNeuralTube^e^ attribution scores were calculated using the CREsted^24^ (v1.3.0; RRID:SCR_026617) function integrated grad, on 1,000 regions with the highest prediction score for that topic according to the model. Next, patterns/motifs were discovered *de novo* using a custom implementation of tfmodisco-lite (v2.0.0; RRID:SCR_026709). First seqlets were called using the tfmodisco-lite function extract_seqlets using the attribution scores as input and using parameters sliding_window_size = 15, flank = 5, suppress = 12, target_fdr = 0.2, min_passing_windows_frac = 0.03, max_passing_windows_frac = 0.3 and weak_threshold_for_counting_sig = 0.8. Next, the attribution score of each seqlet was extracted based on the seqlet coordinates and negative and positive seqlets were defined based on the sum of the attribution score without the flank filtering out seqlets for which the absolute value of the sum of the attribution score was lower than or equal to the threshold value obtained using the tfmodisco-lite function extract_seqlets. Finally, separately for positive and negative seqlets 1,000 seqlets were sampled and used in the tfmodisco-lite function seqlet to patterns using parameters min_overlap_while_sliding = 0.7, nearest_neighbors_to_compute = 500, affmat_correlation_threshold = 0.15, tsne_perplexity = 10, n_leiden_iterations = -1, n_leiden_runs = 50, frac_support_to_trim_to = 0.2, min_num_to_trim_to = 30, trim_to_window_size = 20, initial_flank_to_add = 5, prob_and_per_track_sim_merge_thresholds = [(0.8,0.8), (0.5, 0.85), (0.2, 0.9)], prob_and_per_track_sim_dealbreaker_thresholds = [(0.4, 0.75), (0.2,0.8), (0.1, 0.85), (0.0,0.9)], subcluster_perplexity = 50, merging_max_seqlets_subsample = 300, final_min cluster_size = 20, min_ic_in_window = 0.6, min_ic_window_size = 6 and ppm_pseudocount = 0.001.

### Clustering of *de novo* TF-MoDISCo and known- and enriched motifs

A pairwise similarity matrix between all enriched motifs (see Motif enrichment analysis) and TF-MoDISCo *de novo* motifs (see *De novo* motif discovery using TF-MoDISCo) was calculated using TomTom^107^ implemented in tangermeme^108^ (v0.4.0; RRID:SCR_026620). The Scanpy^91^ (v1.11.0; RRID: SCR_018139) functions pca, neighbors, tsne and leiden were used to generate a tSNE-embedding of the motifs based on 50 principle components and to cluster the motifs at a leiden resolution of 2.

### TF-MINDI analysis on human embryo and human neural tube organoid data

For all organoid and embryo topics using respectively DeepNeuralTube° and DeepNeuralTube^e^ attribution scores were calculated using the CREsted^24^ (v1.3.0; RRID:SCR_026617) function integrated grad, on 1,000 regions with the highest prediction score for that topic according to the model. Seqlets were called using the TF-MINDI (v1.1.0; RRID:SCR_027436) function tm.pp.extract_seqlets using threshold = 0.05 and additional_flanks = 10 and a similarity matrix was calculated containing the motif similarity of each seqlet to 3,989 known motifs (https://github.com/aertslab/TF-MINDI/blob/main/paper/sampled_motifs.txt) using the TF-MINDI function tm.pp.calculate_motif_similarity, using default arguments. Finally, an AnnData^75^ (v0.12.2; RRID:SCR_018209) was generated using the TF-MINDI function tm.pp.create_seqlet_adata. Resulting pattern logos were visually inspected and Sox dimer, Coordinator and NKX motifs were manually annotated.

### TF-MINDI Code table generation

Two matrices were generated: one containing the average number of instances per TF-family and region and another containing the average instance-contribution score per TF-family and region. Both matrices were visualized using a dotplot where the dotsize represents the average number of instances and the color the average contribution score.

### Calculating overlap between TF-MINDI instances from organoid and embryo

TF-MINDI (v1.1.0; RRID:SCR_027436) instances were saved as bed files, containing their genomic position, per TF-family. This was done separately for organoid and embryo results. The genomic overlap between pairs of bed files (one from organoid and the other from embryo) were calculated using the bedtools^109^ (v2.31.0; RRID:SCR_006646) command intersect and the jaccard index was calculated for each pair.

### Linking TF-MINDI patterns to TFs

A matrix containing the average expression per accessibility topic was calculated based on the binarized cell topic probabilities. Next, for each TF-MINDI seqlet the best motif match was obtained and for this match a list of candidate transcription factors was obtained using the “direct_annot” and “Orhtology_annot” field from the motif-to-TF annotation database (obtained through the TF-MINDI (v1.1.0; RRID:SCR_027436) function tm.fetch_motif_annotations and tm.load_motif_annotations). Next, for each TF-family the Pearson correlation coefficient was calculated between the average number of TF-family instances per region (see TF-MINDI Code table generation) and the average expression of the candidate TFs for that TF-family per accessibility topic, using the Scipy^94^ (v1.9.3; RRID:SCR_008058) stats.pearonsr. Only TFs with a correlation coefficient greater than 0.5 were retained.

### TF-MINDI analysis on human embryo and human neural tube organoid data

A CREsted (v1.3.0; RRID:SCR_026617) dilated cnn model was trained on all 134 clusters defined in Mannens *et al*., 2024^57^. First, bigwigs were normalized using the CREsted normalize_peaks function based on the top 3% of peaks. The dilated cnn model had an input sequence length of 2114, 8 convolutional layers with batch normalization, a fully connected dense layer of dimension 512 and a dense output layer of dimension 134 to predict peak height based on genomic sequence. For the activation function RELU was used for the convolutional layers, with a kernel size of 3 and a dropout of 0.1. Convolutional layers were l2-normalized using an epsilon value of 1e-5, the Adam optimizer had a learning rate of 1e-3 with CosineMSELog as the loss function and a batch size of 256. Tensorflow (v2.16.3; RRID:SCR_016345) was used to build and train the model. Both the forward and reverse complement sequences were used for training and validation and testing was done using 10% of the data each. The training was stopped after 15 epochs. The model was then finetuned on highly variable peaks (Gini index 1 standard deviation above the mean) with a learning rate of 1e-4 and stopped after 24 epochs.

The attribution scores for the 1,000 regions with the highest Gini score was calculated using the CREsted integrated grad function, after which the central 500 bp were excised to match the region size of the topic models.

Seqlets were called with the same settings as were used for the DeepNeuralTube^e^ and DeepNeuralTube° (threshold = 0.05, additional_flanks = 10) and motif similarity to the 3,989 known motifs was calculated. The resulting anndata object was concatenated with the DeepNeuralTube^e^ and DeepNeuralTube° objects using the TF-MINDI tm.concat function.

The Scanpy91 (v1.11.5; RRID: SCR_018139) functions pca, neighbors, tsne and leiden were used to generate a tSNE-embedding of the motifs based on 50 principle components and to cluster the motifs at a leiden resolution of 3.

### TWIST1 ChIP-seq enrichment of human organoid and human embryo TF-MINDI instances

A TWIST1 ChIP-seq BigWig was downloaded from ChIP-atlas^110^ (SRX0052931) and for each TF-MINDI seqlet ChIP-seq signal was extracted using the pyBigWig^111^ (v0.3.24; RRID:SCR_024807) stats function. Finally, ROC-curves were calculated using the Scikit-learn^105^ (v1.5.2; RRID:SCR_002577) function roc_curve using the maximum ChIP-seq signal across seqlets per region as y_score and a binary vector indicating whether each region contains a seqlet annotated as coordinator as y_pred. Motif enrichment based ROC-curves were calculated similarly using a binary vector indicating whether the region is in the leading edge of any motif annotated to TWIST1 as y_pred. To calculate the recovery of TWIST1 ChIP-seq peaks the following procedure was used. First TF-MINDI was applied to facial mesenchyme regions (see below: TF-MINDI analysis on organoid facial mesenchyme regions) and a bed file was generated from the instances annotated as coordinator. Next, a bed file containing TWIST1 ChIP-seq peaks was downloaded from ChIP-atlas (SRX0052931) and the recovery of each peak was calculated using the bedtools^109^ (v2.31.0; RRID:SCR_006646) command intersect.

### TF-MINDI analysis on human peripheral blood mononuclear cell data

Attribution scores were calculated using the CREsted^24^ (v1.3.0; RRID:SCR_026617) function integrated grad, on 2,000 screen v4 regions^112^ with the highest prediction score for each class of deepPBMC^24^. Next, seqlets were called using the TF-MINDI (v1.1.0; RRID:SCR_027436) function tm.pp.extract_seqlets on the 500 bp center of each region using parameters threshold = 0.05 and additional_flanks = 10 and a similarity matrix was calculated containing the motif similarity of each seqlet to 3,989 known motifs (https://github.com/aertslab/TF-MINDI/blob/main/paper/sampled_motifs.txt) using the TF-MINDI function tm.pp.calculate_motif_similarity, using default arguments. Finally, an AnnData^75^ (v0.12.2; RRID:SCR_018209) was generated using the TF-MINDI function tm.pp.create_seqlet_adata.

### Transcription factor ChIP-seq enrichment of human peripheral blood mononuclear cell TF-MINDI instances

ChIP-seq BigWigs were downloaded from Encode^113–115^ for the transcription factors: EBF1 (ENCFF810XRY), IRF3 (ENCFF906RAB), IRF4 (ENCFF167KPF), IRF5 (ENCFF105OXO), PAX5 (ENCFF914QGY), ETS1 (ENCFF686FAA), STAT1 (ENCFF840IFX), TCF12 (ENCFF203JRX); and for each TF-MINDI seqlet ChIP-seq signal was extracted using the pyBigWig^111^ (v0.3.24; RRID:SCR_024807) stats function. Finally, for each transcription factor, ROC-curves were calculated using the Scikit-learn^105^ (v1.5.2; RRID:SCR_002577) function roc_curve using the ChIP-seq signal per seqlet as y_score and a binary vector indicating whether each seqlet is annotated to the TF-family of the TF as y_pred. For this the following mapping was used: EBF1: EBF1, ETS1: Ets, IRF3: Irf, IRF4: Irf, IRF5: Irf, PAX5: Paired box, TCF12: bHLH. Motif enrichment based ROC-curves were calculated as follows: First, for each TF, regions in the leading edge of motifs annotated to that TF were obtained and saved as bed files. Second, bed files were generated for TF-MINDI seqlets. Third, a binary vector indicating whether a seqlet lies within a TF-leading edge region was produced using the bedtools^109^ (v2.31.0; RRID:SCR_006646) command intersect using the -c option. Finally, this binary vector was used as the y_pred parameter in the Scikit-learn function roc_curve.

### TF-MINDI analysis on organoid facial mesenchyme regions

Attribution scores were calculated using the CREsted^24^ (v1.3.0; RRID:SCR_026617) function integrated grad on all regions of organoid accessibility topic 57 (after performing topic binarization) using DeepNeuralTube°. Next, seqlets were called using the TF-MINDI (v1.1.0; RRID:SCR_027436) function tm.pp.extract_seqlets using threshold = 0.05 and a similarity matrix was calculated containing the motif similarity of each seqlet to 3,989 known motifs (https://github.com/aertslab/TF-MINDI/blob/main/paper/sampled_motifs.txt) using the TF-MINDI function tm.pp.calculate_motif_similarity, chunk_size = 10,000. Coordinator seqlets were detected by centering on either bHLH or Homeodomain instances, obtaining attribution scores within a 10 bp window up- and downstream of the instances and detecting a secondary instance that passes an average z-score threshold of 0.1 or 0.4 when centering on respectively bHLH or Homeodomain instances. If a secondary instance was detected, the seqlet was extended and original instances that overlap with the seqlet were detected using the ncls (v0.0.63; RRID:SCR_027849) package and removed to avoid duplicate seqlets. Finally, an AnnData^75^ (v0.12.2; RRID:SCR_018209) was generated using the TF-MINDI function tm.pp.create_seqlet_adata.

### Calculating TWIST1 motif score

The motif TWST1_HUMAN.H11MO.0.A from the Hocomoc motif collection v11^116^ was used to score all organoid accessibility topic 57 regions. For this, the genomic regions were onehot-encoded and the position weight matrix was converted to a position probability matrix by dividing by the sum of values for each position. Next, the PWM score was generated across a sliding window with window size equal to the size of the PWM and using the NumPy^117^ (v2.0.1; RRID:SCR_008633) lib.stride_tricks.sliding_window_view and einsum functions. The same procedure was used to score all TF-MINDI facial mesenchyme seqlets. However, in case the size of the seqlet was smaller than the position weight matrix it was zero-padded.

### Design of synthetic enhancers

10,000 random DNA sequences were generated that follow the local nucleotide distribution of consensus peaks. For this, the fraction of each nucleotide per position across all consensus peaks was calculated and new DNA sequences were generated by randomly sampling nucleotides according to this distribution. The middle 160 bp of each random DNA sequence was used to design enhancers. First, up to two coordinator instances were implanted at varying motif scores, ensuring a minimal distance of 5 bp between subsequent implantations in the same sequence. Coordinator instances were sampled from the TF-MINDI seqlets across 20 equal size bins according to the TWIST1 motif score. Next, up to five additional motifs were inserted in the same DNA sequences, ensuring a minimal distance of 3 bp between already inserted motifs. These additional motifs were sampled from TF-MINDI seqlets of manually selected patterns (showing good quality motif logos). For each pattern, only the top 5% most frequently occurring seqlets were considered.

### Luciferase reporter assay of synthetic enhancers

#### Selecting sequences to test

For selecting negative and positive control sequences, the prediction score of DeepNeuralTube° and DeepNeuralTube^e^ was calculated on all organoid consensus peaks for the facial mesenchyme class of both organoid (accessibility topic 57) and embryo (accessibility topic 100). Two positive control sequences were sampled randomly from consensus peaks that had a prediction score greater than 0.85 for both models and two negative control sequences were sampled randomly from consensus peaks that had a prediction score smaller than 0.05 for both models. Three sequences in which we implanted two coordinator motifs, of the highest motif score bin, were randomly sampled without considering the model’s prediction score. For these sequences, we also selected the version where only a single coordinator instance (of the highest affinity bin) was inserted and the version where a single coordinator instance (of the highest affinity bin) was inserted along with five additional motifs.

#### Cloning enhancers in luciferase enhancer-reporter plasmid

DNA sequences were ordered from Twist Bioscience (RRID: SCR 025817) precloned in the pTwist ENTR vector and sub-cloned in the pGL4.23-GW (Promega; RRID:Addgene_60323) luciferase reporter vector through Gateway LR recombination reaction (Invitrogen). Briefly, a reaction mixture was generated containing 150 ng pGL4.23-GW plasmid DNA, 20ng enhancer plasmid DNA, 1x Gateway LR Clonase enzyme (ThermoFisher, 11791020) in 1µl TE buffer which was incubated for 1h at 25°C after which Proteinase K (ThermoFisher, 11791020) 0.33 µg/µl was added followed by 10 min incubation at 37°C. Next, 2.5µl of the reaction mix was transformed into 25µl Stellar chemically competent bacteria (Takara, 636763). By thawing bacteria on ice for 15 min, add reaction mixture, keep 30 min on ice, 45 sec heat shock at 42°C and 5 min ice. Bacteria were incubated for 1h at 37°C in 250µl SOC medium (Takara, 636763) and plated on Carbencilin plates and incubated overnight at 37°C. Next, single colonies were picked and grown in 8ml LB medium, and plasmids were isolated using Qiagen miniprep kit (Qiagen, 27104) and sequenced using Sanger sequencing to confirm correct insertion in the destination plasmid.

#### Transfection and luciferase assay

Facial mesenchyme cells were seeded in a 24 well tissue culture plate, coated with fibronectin and in 200 µl long term maintenance medium, to reach 80% confluency the next day. The next day, medium was changed to 300 µl long term maintenance medium. A mixture containing 0.2 µg/µl enhancer-reporter plasmid and 1 ng/µl pRL-TK Renilla plasmid (Promega; RRID:Addgene_27163) and 6% (V/V) Fugene HD Transfection Reagent (Promega, E2311) in Opti-MEM (ThermoFisher, 31985062). After incubating the mixture for 10 min at room temperature 30 µl is added to each well of the 24 well plate dropwise and cells are incubated overnight. Finally, luciferase activity was measured through the Dual-Luciferase Reporter Assay System (Promega), following the manufacturer’s protocol. Cells were washed once using PBS and lysed with 1x passive lysis buffer for 15 min gently shaking the plate. A total 20 µl of the lysate was transferred in duplicate in a well of an OptiPlate-96 HB (PerkinElmer) and 100 µl of luciferase assay reagent II was added in each well. Luciferase-generated luminescence was measured on a Victor X luminometer (PerkinElmer). A total of 100 µl of the Stop & Glo Reagent was added to each well and the luminescence was measured again to record Renilla activity. Luciferase activity was estimated by calculating the ratio luciferase/Renilla; this value was normalized by the ratio calculated on blank wells containing only reagents.

### TF-MINDI topic modeling

A count table containing the number of instances per TF-MINDI cluster across consensus peaks was generated for organoid and embryo data. In case a region occurred for multiple topics, the region with the highest black box prediction score was taken. Furthermore, only regions with a minimal prediction score of 0.4 were considered. Topic modeling using the python lda (RRID:SCR_027850) was performed on this matrix using 2,000 iterations, eta = 0.1, alpha = 0.04 and a model with 25 topics for the organoid data and a model with 20 topics for the embryo data.

### Zebrafish development analysis

#### Data and quality control

scATAC-seq fragment files and cell type annotation metadata were downloaded from https://zebrahub.sf.czbiohub.org for the following samples: TDR118, TDR119, TDF124, TDR125, TDR126, TDR127 and TDR128. Quality metrics were calculated using the pycisTopic^52^ (v1.0.3.dev20+g8955c76, RRID:SCR_026618) qc command and cells were filtered based on the following thresholds:

● TDR118: 3000 minimum number of unique fragments, 8 minimal TSS enrichment, 0.4 minimal fraction of reads in peaks
● TDR119: 2000 minimum number of unique fragments, 8 minimal TSS enrichment, 0.4 minimal fraction of reads in peaks
● TDR124: 500 minimum number of unique fragments, 8 minimal TSS enrichment, 0.4 minimal fraction of reads in peaks
● TDR125: 3000 minimum number of unique fragments, 8 minimal TSS enrichment, 0.4 minimal fraction of reads in peaks
● TDR126: 2500 minimum number of unique fragments, 8 minimal TSS enrichment, 0.4 minimal fraction of reads in peaks
● TDR127: 2400 minimum number of unique fragments, 8 minimal TSS enrichment, 0.4 minimal fraction of reads in peaks
● TDR128: 2200minimum number of unique fragments, 8 minimal TSS enrichment, 0.4 minimal fraction of reads in peaks Resulting in 104,129 high-quality cells

#### Sequence-to-function training

A topic classification model was trained using CREsted^24^ (v1.5.0; RRID:SCR_026617). A validation and test fraction of 10 % was used, shift augmentation of 3 bps, reverse complement augmentation, batch size of 256 and input size 500 bps. The default deeptopic cnn was used as architecture and default training configurations.

### TF-MINDI analysis on zebrafish development data

For all zebrafish topics, attribution scores were calculated using the CREsted^24^ (v1.3.0; RRID:SCR_026617) function integrated grad, on 1,000 regions with the highest prediction score for that topic. Seqlets were called using the TF-MINDI (v1.1.0; RRID:SCR_027436) function tm.pp.extract_seqlets using threshold = 0.05 and additional_flanks = 10 and a similarity matrix was calculated containing the motif similarity of each seqlet to 3,989 known motifs (https://github.com/aertslab/TF-MINDI/blob/main/paper/sampled_motifs.txt) using the TF-MINDI function tm.pp.calculate_motif_similarity, using default arguments. Finally, an AnnData^75^ (v0.12.2; RRID:SCR_018209) was generated using the TF-MINDI function tm.pp.create_seqlet_adata.

### Human and zebrafish seqlet integration and scoring of zebrafish regions using topic modeling

The AnnData^75^ (v0.12.2; RRID:SCR_018209) objects containing TF-MINDI results on the organoid, embryo and zebrafish data were combined using the anndata concat function. Next, zebrafish TF-MINDI seqlets were annotated to both organoid and embryo clusters by performing leiden clustering on the combined TF-MINDI object using resolution, TF-MINDI (v1.1.0; RRID:SCR_027436) function tm.tl.cluster_seqlets and annotating the clusters based on a majority vote of organoid or embryo seqlet-cluster annotations. Next, a count table was generated containing the number of seqlet instances per organoid or embryo cluster across zebrafish regions and each region was scored for the organoid or embryo topic model using the lda (RRID:SCR_027850) transform method.

### Chicken embryo enhancer reporter assay

#### Enhancer cloning

Enhancer candidates were ordered from Twist Bioscience (RRID: SCR 025817) with 5’ GGTACCGAGCTCGAG and 3‘ GGGCTCGAGATCTGC adaptors. Plasmid pTK BsmBI LacZ Citrine Nanotag 24^118^ (RRID:Addgene_130522) was linearised using forward primer 5‘-ggAACTCGAGCTCGGTACCT-3‘ and reverse primer 5‘-GGGCTCGAGATCTGCGATC-3‘ using PCR. A PCR mixture of KAPA HiFi hot start ready mix (Roche, 07958935001 50 % V/V), 0.8 μg/μl plasmid and 0.3 μM of forward and reverse primer and PCR program: 95°C for 3 minutes, 15 cycles of: 98°C for 20 seconds, 58 °C for 15 seconds and 72 ° C for 2.5 minutes, followed by 72 °C for 5 minutes was used. Linearised plasmids were purified using QIAquick PCR purification kit (Qiagen, 28104), followed by DpnI digestion for 1 hour at 37°C and gel extraction of 4,5 kb band using QIAquick Gel Extraction Kit (Qiagen, 28704). Enhancer candidates were cloned into the target vector using an NEBuilder (New England Biolabs, E5520S) reaction with a linearised plasmid to insert ratio of 1:2 (45 minutes at 50°C. Next, 2.5 μl of the reaction mixture was transformed into 25 μl Stellar Competent Cells (Takara Bio, 636763). Cells were thawed 15 minutes on ice, plasmid + cell mixture was kept for an additional 30 minutes on ice followed by 45 second heat shot at 42°C and 5 minutes on ice. Cells were incubated for 1 hour at 37°C, single colonies were then grown on Carbencillin plates over night. Finally, plasmids were purified using NucleoBond Xtra Maxi Plus Kit (Macherey-Nagel, 740416.50) and eluded at a target concentration of 4 μg per μl.

#### Injection control cloning

Plasmid pCI H2B-RFP^118^ (RRID:Addgene_92398) was ordered from Addgene, transformed into Stellar Competent Cells (Takara Bio, 636763). Cells were thawed 15 minutes on ice, plasmid + cell mixture was kept for an additional 30 minutes on ice followed by 45 second heat shot at 42 °C and 5 minutes on ice. Cells were incubated for 1 hour at 37 °C, single colonies were then grown on Carbenicillin plates overnight. Finally, plasmids were purified using NucleoBond Xtra Maxi Plus Kit (Macherey-Nagel, 740416.50) and eluded at a target concentration of 4 μg/μl.

#### Ex ovo enhancer-reporter injection

Preparation of chicken embryos and injection was performed as described in Williams and Sauka-Spengler 2021^119^. Eggs were incubated as described above (see Chicken embryos). A hole was created in the egg shell using the back of forceps, thick albumin was discarded and thin albumin was collected. When most albumin was removed, the egg shell was cut open and the yolk turned so that embryo faces upwards. More albumin was removed from the yolk by stroking the yolk using serrated forceps, moving away from (and not touching) the embryo. A Whatman filter paper with perforated hole was placed on top of the embryo and yolk and whatman paper was cut using dissection scissors and embryo was removed and placed in Ringer solution (NaCl_2_ 62.5 μM; KCl 5 μM; CaCl_2_ · 2H_2_O 1.5 μM; NaHPO_4_ · 7H_2_O 0.75 μM; KH_2_PO_4_ 0.15 μM). Embryo was placed in custom electroporation cuvette (see Williams and Sauka-Spengler 2021^120^), enhancer-reporter plasmid mixture (enhancer-reporter plasmid 2 μg/μl; injection control plasmid 1 μg per μl; food colouring) was injected using micro injector in the cavity between the blastoderm and underlying vitelline membrane and embryo was electroporated (5 pulses of 5V: 100 ms on, 50 ms off). Embryo was grown for 24 hours at 37°C and imaged whole-mount or sectioned and imaged.

#### Chicken embryo cryosectioning

Embryos were fixed in 4 % paraformaldehyde for 1 hour at room temperature followed by three 10 minute washes with PBS. Embryos were transferred to 15 % V/V sucrose in PBS at room temperature until they sank to the bottom. Next, embryos were transferred to a mixture consisting of 7.5 % V/V gelatine and 15 % V/V sucrose in PBS and were incubated overnight at 37°C. In the meantime 20 % V/V gelatine was dissolved in PBS at 60°C and this was kept at 37°C once the gelatine was dissolved. After the overnight incubating, embryos were transferred to the gelatine solution and kept for a minimum of 4 hours at 37 °C. Finally, embryos in gelatine solution were transferred to a cryomold and quickly frozen on an aluminium block cooled by dry ice. Cryosections of 20 μm were made at -60°C and collected on SuperFrost Plus Adhesion slides (Fisher Scientific, 10149870). Sections were stained with DAPI 0.1 mg/ml in PBS; Sigma-Aldrich, 28718-90-3) for 15 minutes, followed by a PBS wash and air dried. Cover slip was mounted using Mowiol 4-88 mounting medium (Sigma-Aldrich, 81381-50G).

## Data availability

Raw sequencing data, scATAC-seq fragment files, scRNA-seq count matrix, cell metadata, consensus peaks, and cell- and region topic probabilities for the human neural tube organoids are available on Gene Expression Omnibus (GEO) using accession number https://www.ncbi.nlm.nih.gov/geo/query/acc.cgi?acc=GSE295337. Raw sequencing data of the human embryo is available through the European Genome Phenome Archive using accession numbers: EGAD50000001501 (RNA) and EGAD50000001502 (ATAC). scATAC-seq fragment files, scRNA-seq count matrix, cell metadata, consensus peaks and cell-and region topic probabilities of the human embryos are available through Zenodo (https://doi.org/10.5281/zenodo.15281826). DeepNeuralTube° and DeepNeuralTube^e^ are available via Zenodo (https://doi.org/10.5281/zenodo.15280773). TF-MInDi bed files are available via Zenodo (https://doi.org/10.5281/zenodo.16039414). Data can be interactively explored using the https://scope.aertslab.org/#/Neural Tube/*/welcome. TWIST1 ChIP-seq data in facial mesenchyme cells was downloaded fromChIP-atlas (SRX0052931) and EBF1 (ENCFF810XRY), IRF3 (ENCFF906RAB), IRF4 (ENCFF167KPF), IRF5 (ENCFF105OXO), PAX5 (ENCFF914QGY), ETS1 (ENCFF686FAA), STAT1 (ENCFF840IFX) and TCF12 (ENCFF203JRX) in GM12878 cells were downloaded from Encode.

## Code availability

Code for reproducing figures in this manuscript is available on GitHub: https://github.com/SeppeDeWinter/NeuralTube. TF-MInDi code and tutorials are available on: https://github.com/aertslab/TF-MINDI.

## Supplement

### Supplementary note 1

Seqlet similarity matrices derived from deepNeuralTube^e^ and deepNeuralTube° were integrated with a seqlet similarity matrix derived from a peak regression model trained on Mannens et al., (2024)^57^. We observed limited batch effects; separately annotated TF families (on a model to model basis) integrated as expected and several TF families that were not found in the neural tube models, but only in the 6-14 p.c.w embryo, clustered separately (**Fig. S7a-b**). Seqlets specific to 6-14 p.c.w. represent instances of the TF-families: NFI, IRF, MADS box, SMAD and T-box (**Fig. S7a-c**). The NFI family members (*NFIA/B/C*) are expressed in radial glia during the transition to glioblasts and in neurons during differentiation^121^. Instances from the IRF family might reflect *IRF8* expression in microglia^122^. Finally, T-box motif instances are strongly enriched in telencephalic glutamatergic neurons, which coincides with the expression of EOMES, a T-box TF that plays a central role in patterning the forebrain^123^. While the anterior-posterior axis is already established in the 4 p.c.w. neural tube, many of the more fine-grained patterning events only happen in the following weeks.

Conversely, we observe that instances of CUT;HD, Grainyhead and Paired-box are specific to the organoid and 4 p.c.w. embryo. TFs of the Grainyhead family are primarily expressed in the neural crest lineage, which is mostly absent in the later timepoints. Paired-box motifs were present, but relatively sparse in the later embryo despite expression of *PAX2/3/6/7*^57,124^. TFs of the ONECUT (CUT;HD) family are expressed at low levels across the immature neurons in the 6-14 p.c.w. embryo^57,125^, but the model was unable to retrieve the motif.

We next assessed the similarities between floor plate progenitors from the neural tube embryo, organoid and the floor plate cells from the 6-14 p.c.w. embryo, which were on average 11 weeks old. Interestingly, we found that the 11 week old floor plate had transitioned to a state defined primarily by RFX and NFI TF motifs, in contrast to the RFX, TEAD and FOX motifs defining the organoid and early embryo neural tube floor plate progenitors (**Fig. S7f**). This is indicative of a more gliogenic identity as is acquired by radial glial cells during the first trimester^121^.

## Supplementary figures

**Figure S1.**
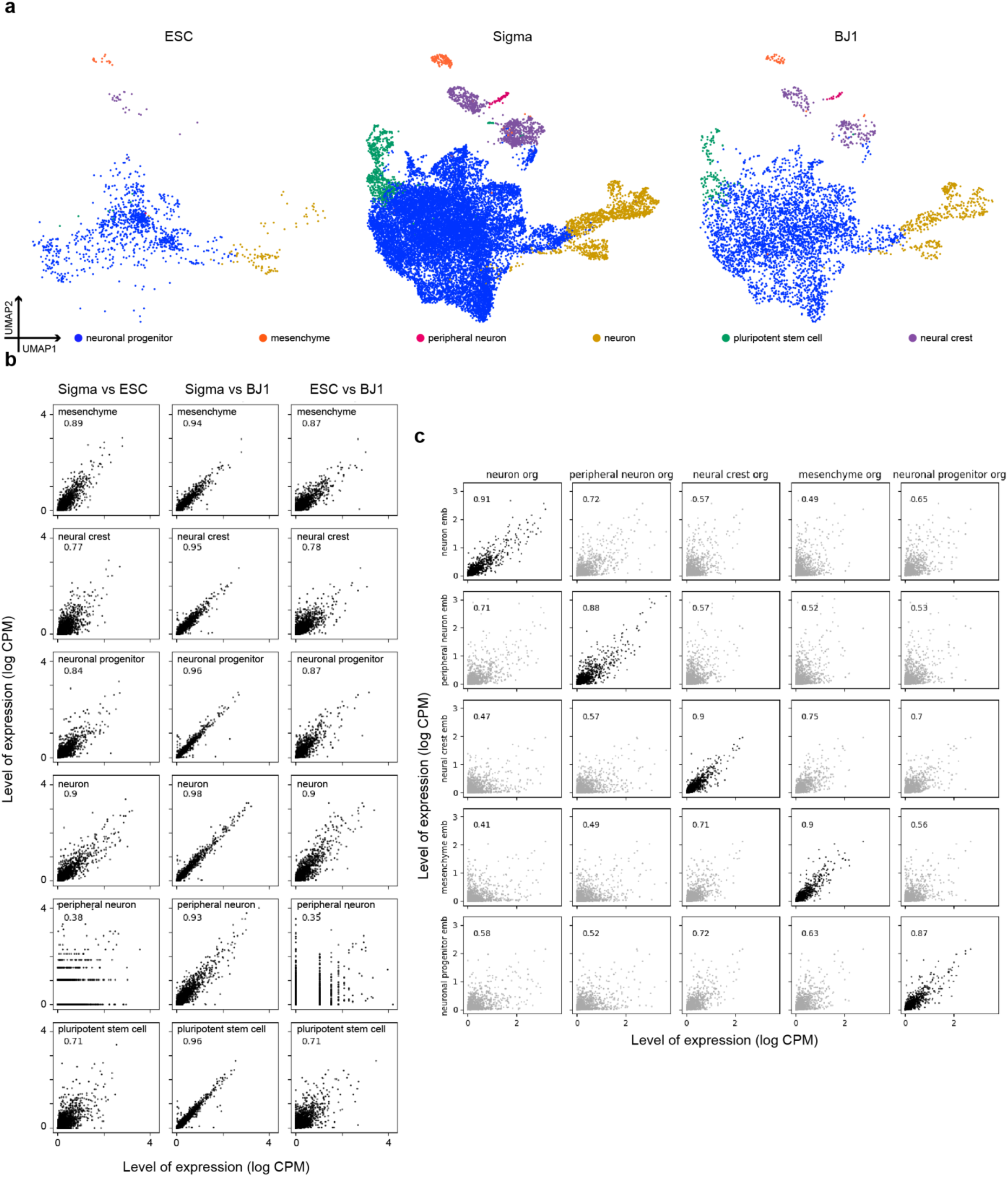
Organoid cell lines and embryo produce similar gene expression profiles. **a**, UMAP based on gene expression profiles of organoid cells colored by cell identity and split by cell line. **b**, scatter plot showing gene expression (of highly variable genes) of the same cell type across cell lines. Number top left indicates Pearson correlation coefficient. **c**, Scatter plot showing pairwise gene expression (of highly variable genes) across cell types and organoid and embryo. Number top left indicates Pearson correlation coefficient.

**Figure S2.**
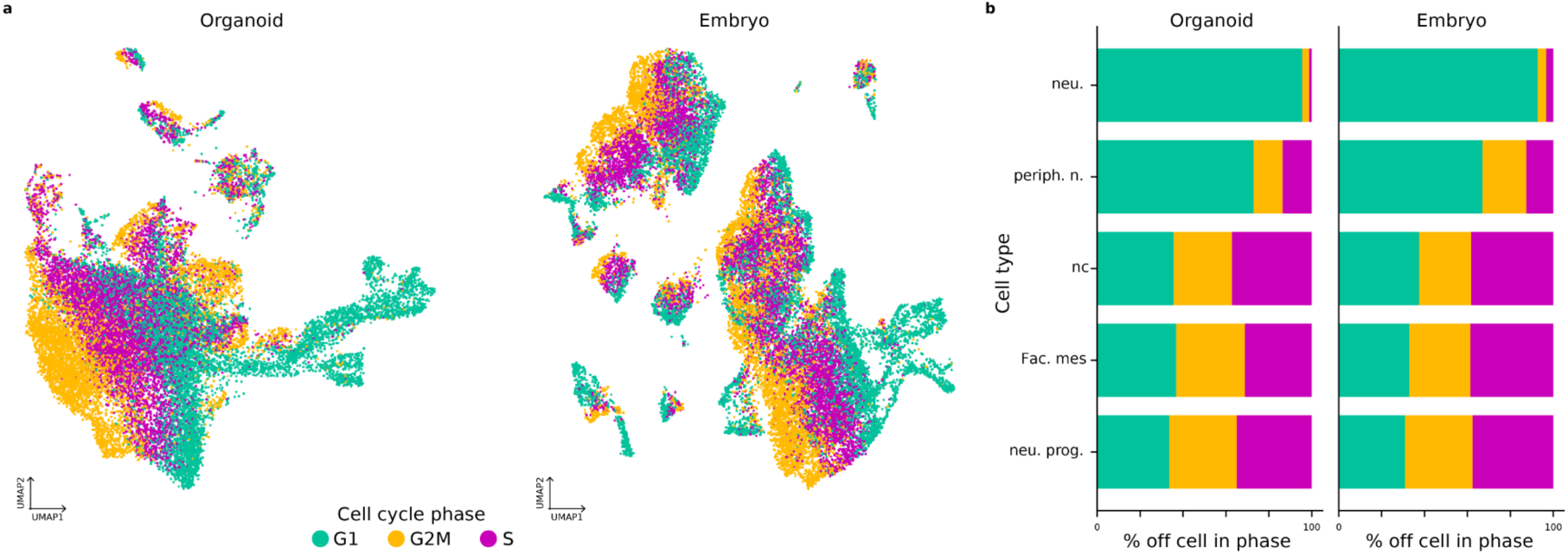
Differentiating neurons are in cell cycle arrest. **a**, UMAP dimensionality reduction of 22,862 organoid cells (left) and 25,785 embryo cells (right) colored by cell cycle phase. **b**, Quantification of percentage of cells per cell type in each phase of the cell cycle. neu.: early differentiating neurons; periph. n; peripheral neurons; NC: neural crest; fac. mes: facial mesenchyme; Neu. prog.: neuronal progenitor.

**Figure S3.**
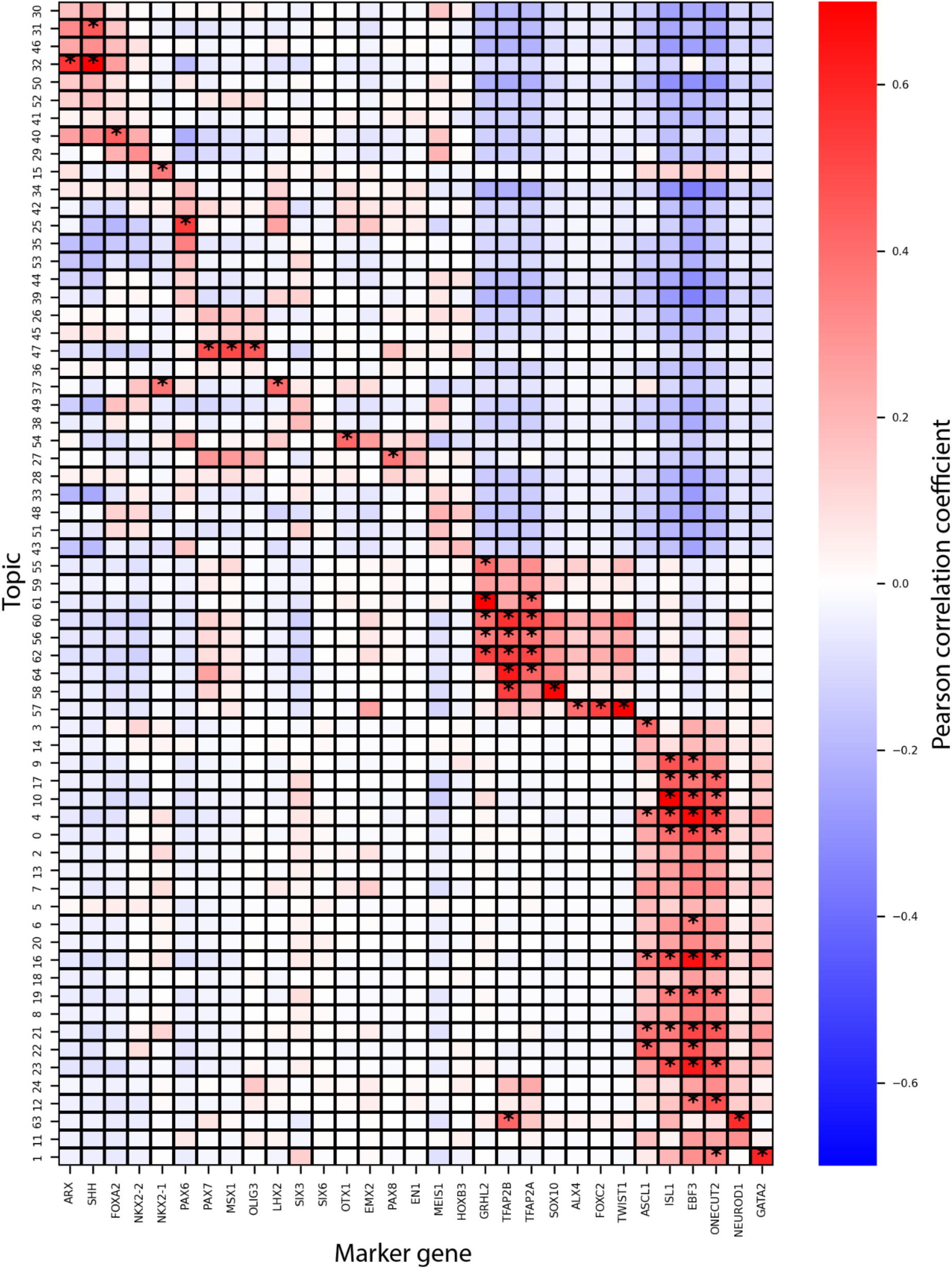
Pearson correlation coefficient between marker gene expression and cell topic probability in organoids. * correlation coefficient > 0.35.

**Figure S4.**
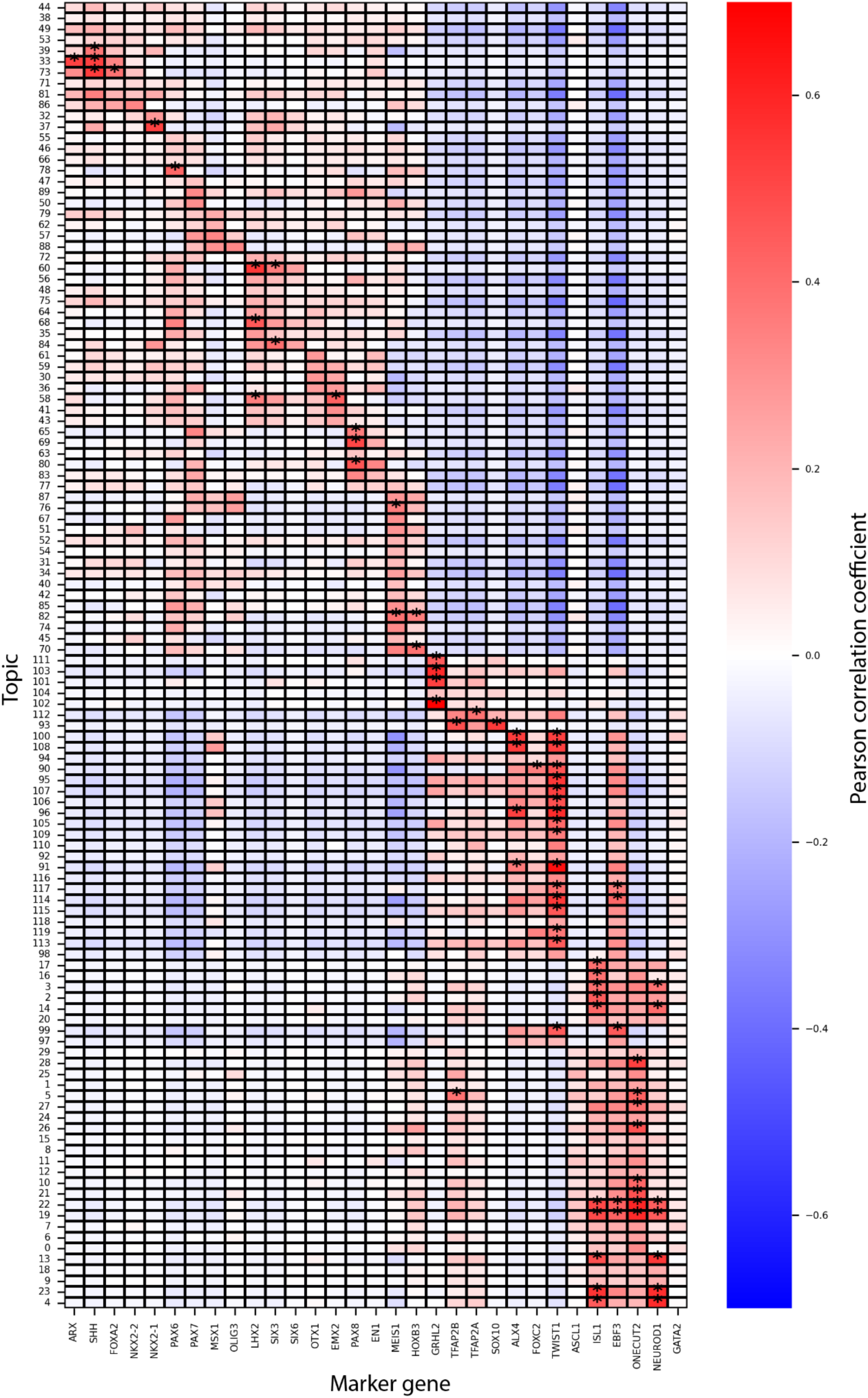
Pearson correlation coefficient between marker gene expression and cell topic probability in embryo. * correlation coefficient > 0.35.

**Figure S5.**
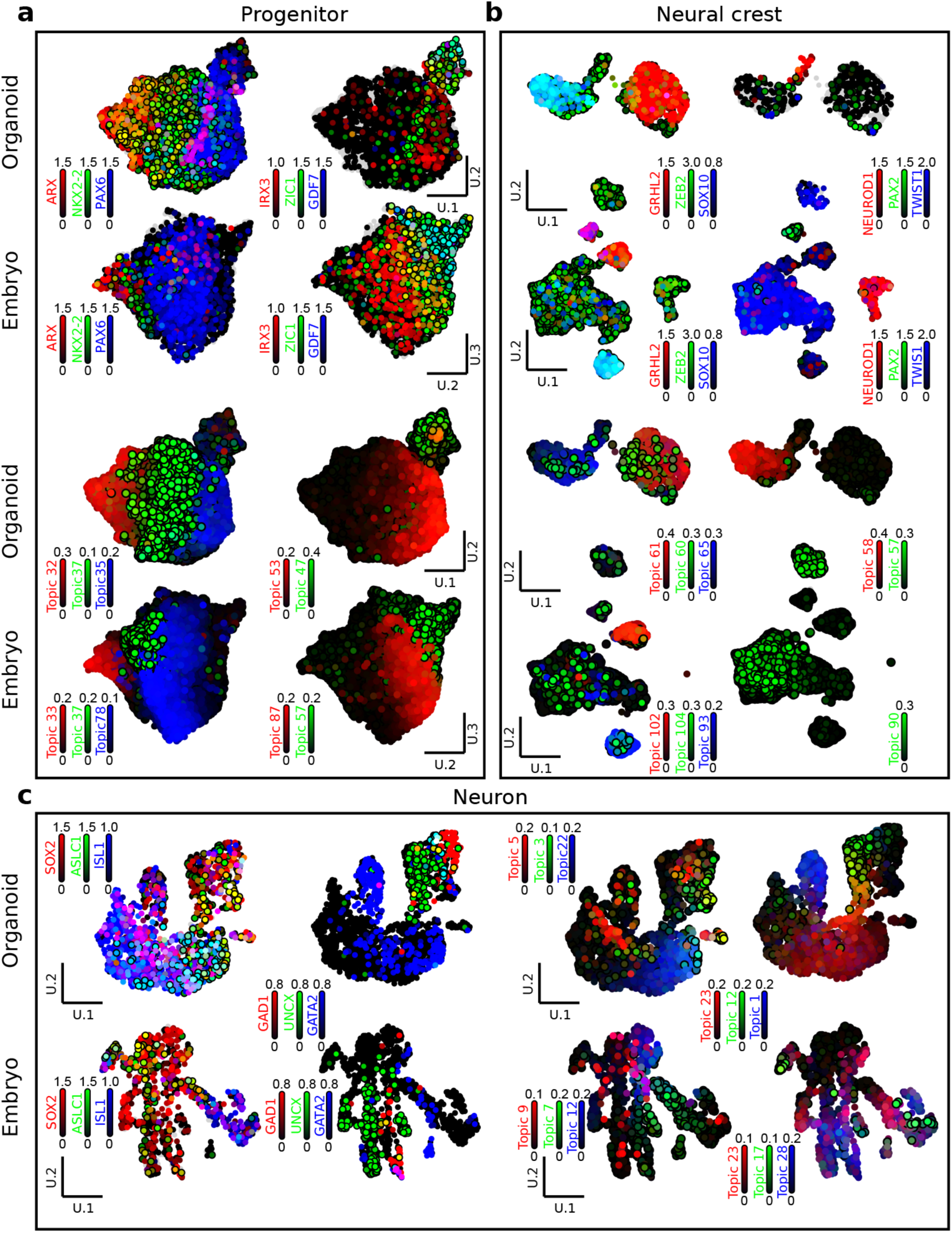
Marker genes and topics in subclustered neuronal progenitors, neural crest and differentiating neurons. UMAP dimensionality reduction of subset of neural progenitors (a), neural crest cells (b) and differentiating neurons (C) colored by the log CPM expression of marker genes and cell topic probability.

**Figure S6.**
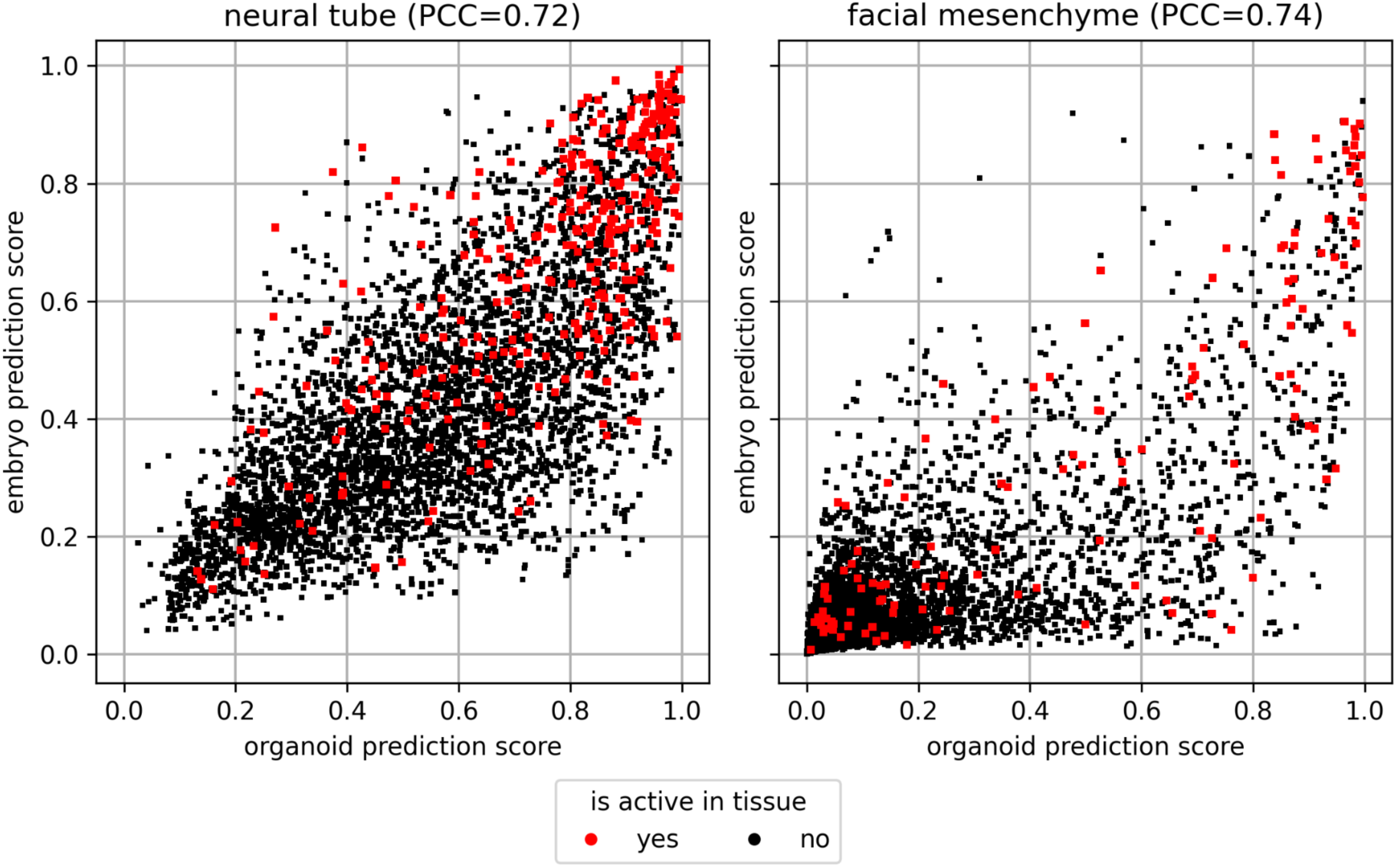
prediction scores of DeepNeuralTube° and DeepNeuralTube^e^ are correlated on VISTA enhancers. Maximum prediction score across neural tube classes for DeepNeuralTube° and DeepNeuralTube^e^ (left) and maximum prediction score across facial mesenchyme classes for DeepNeuralTube° and DeepNeuralTube^e^ (right) of 4,646 sequences from the Vista enhancer browser.

**Figure S7.**
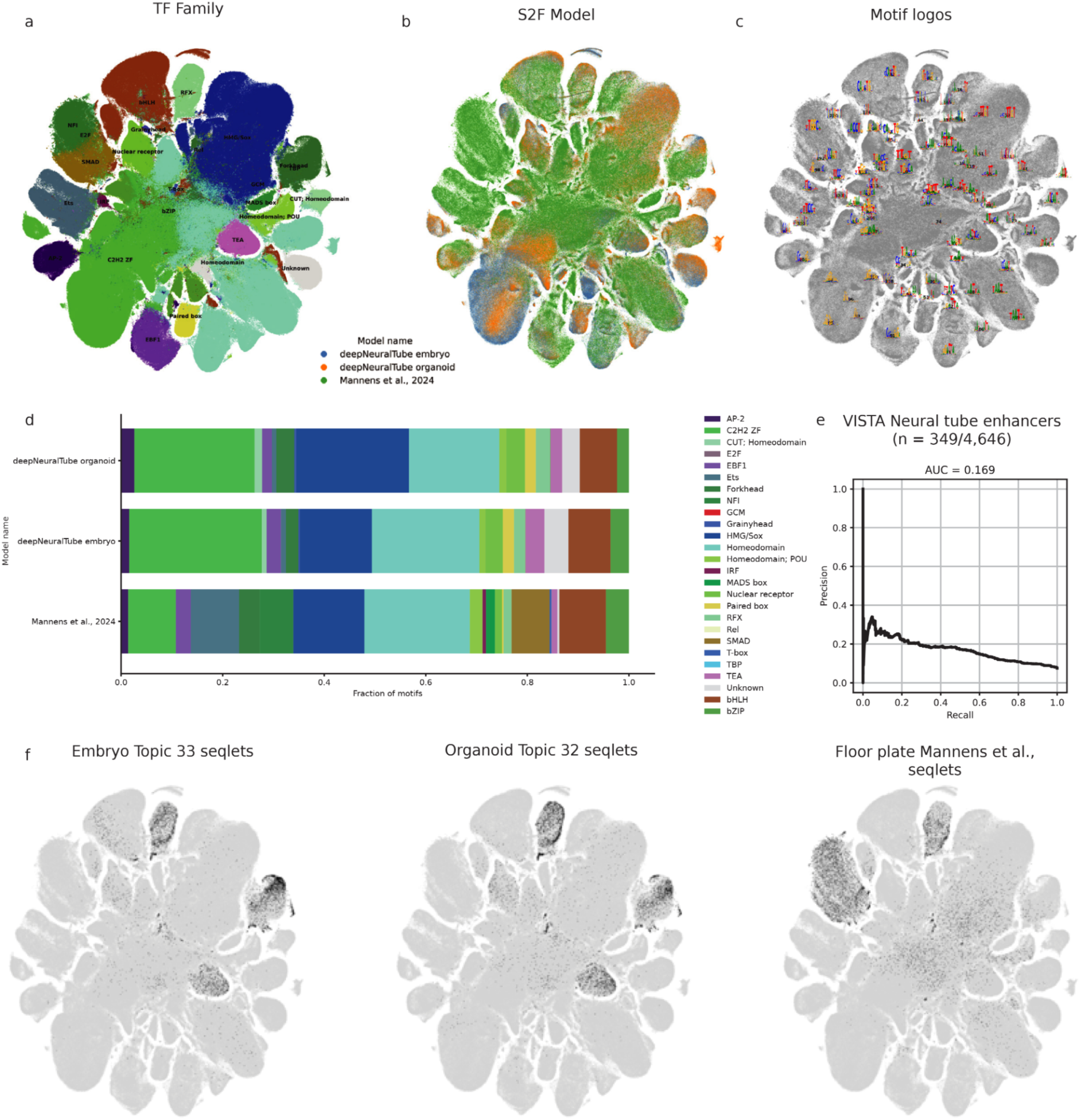
Integration of deepNeuralTube models with a regression model trained on first-trimester whole brain data from Mannens *et al.*, 2024. **a**, Annotation of TF motifs based on similarity to motif database. **b**, Annotation of model used to generate seqlets. **c**, Motlf logos averaged over similarity clusters. **d**, Distribution of TF family annotations across models. Several transcription factor families can only be observed at later developmental timepoints including members of IRF, MADS box, SMAD and T-box. **e.** Precision-recall curve for classifying VISTA neural tube enhancers active in neural vs. non-neural cell types using the first-trimester whole brain regression model. **f**. tSNE maps of seglets derived from floor plate progenitor topics 33 (embryo) and 32 (organoid) contrasted with seqlets from floor plate cells from Mannens *et al.*,

**Figure S8.**
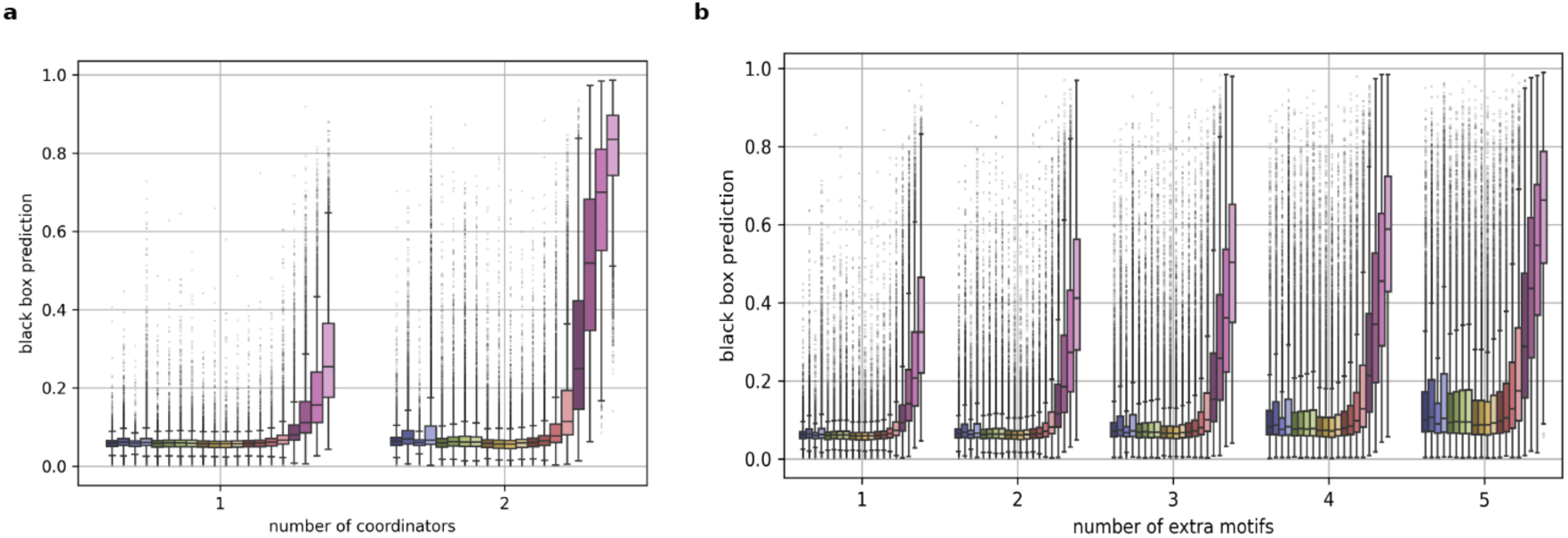
Embryo prediction score on synthetic facial mesenchyme enhancers. DeepNeuralTube^e^ (black box) prediction score after inserting one or two coordinator instances of varying PWM scores (color) into 10,000 random DNA sequence (a), and after adding up to five accessor instances to sequences containing a single coordinator instance at varying affinity (color; b).

**Figure S9.**
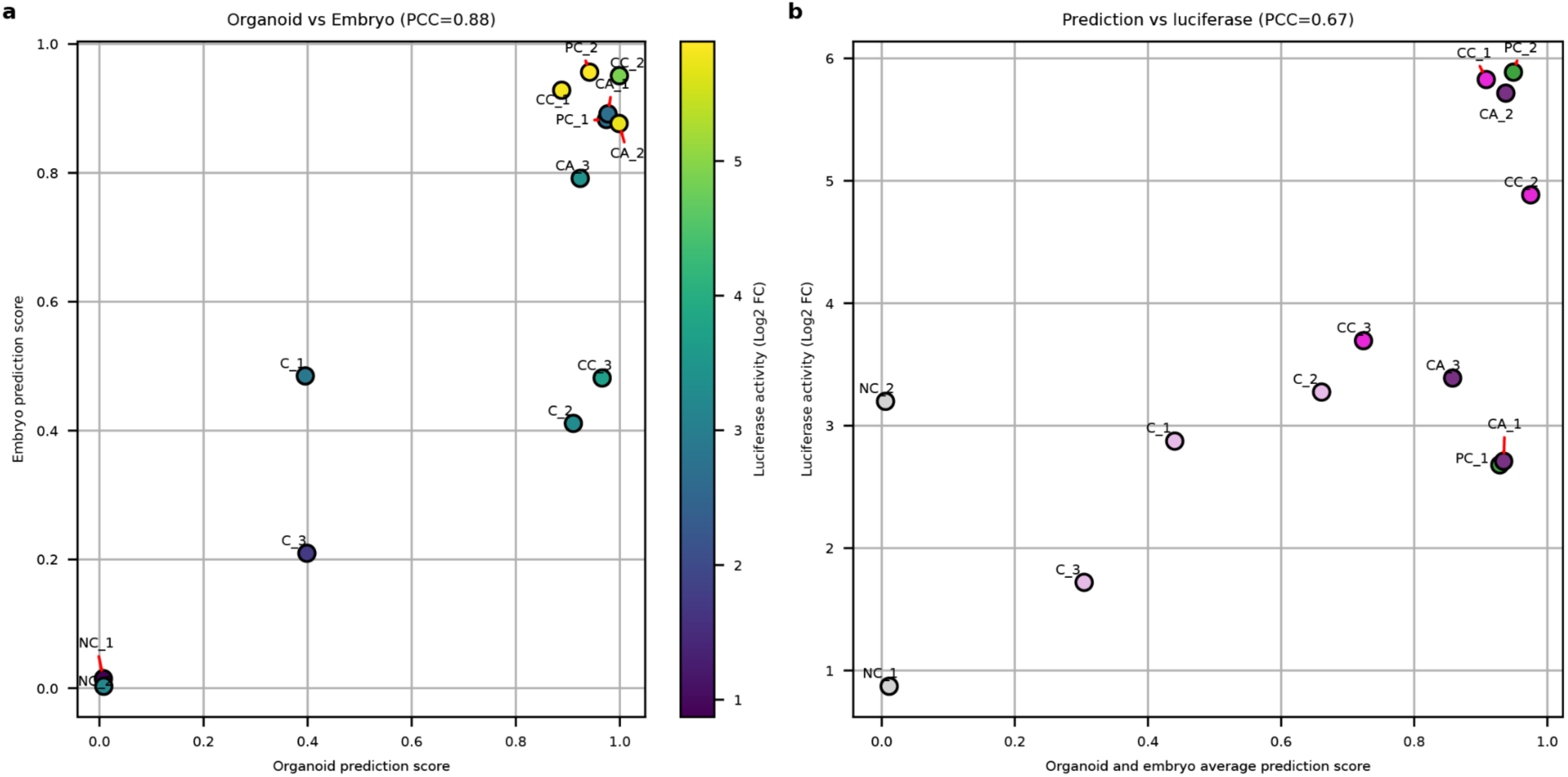
DeepNeuralTube prediction score is correlated to enhancer activity. a. Scatter plot of prediction score of DeepNeuralTube° (for topic 57) and DeepNeuralTube^e^ (for topic 100) colored by luciferase activity. b, Scatter plot of average prediction score of DeepNeuralTube° (for topic 57) and DeepNeuralTube^e^ (for topic 100) and luciferase activity.

**Figure S10.**
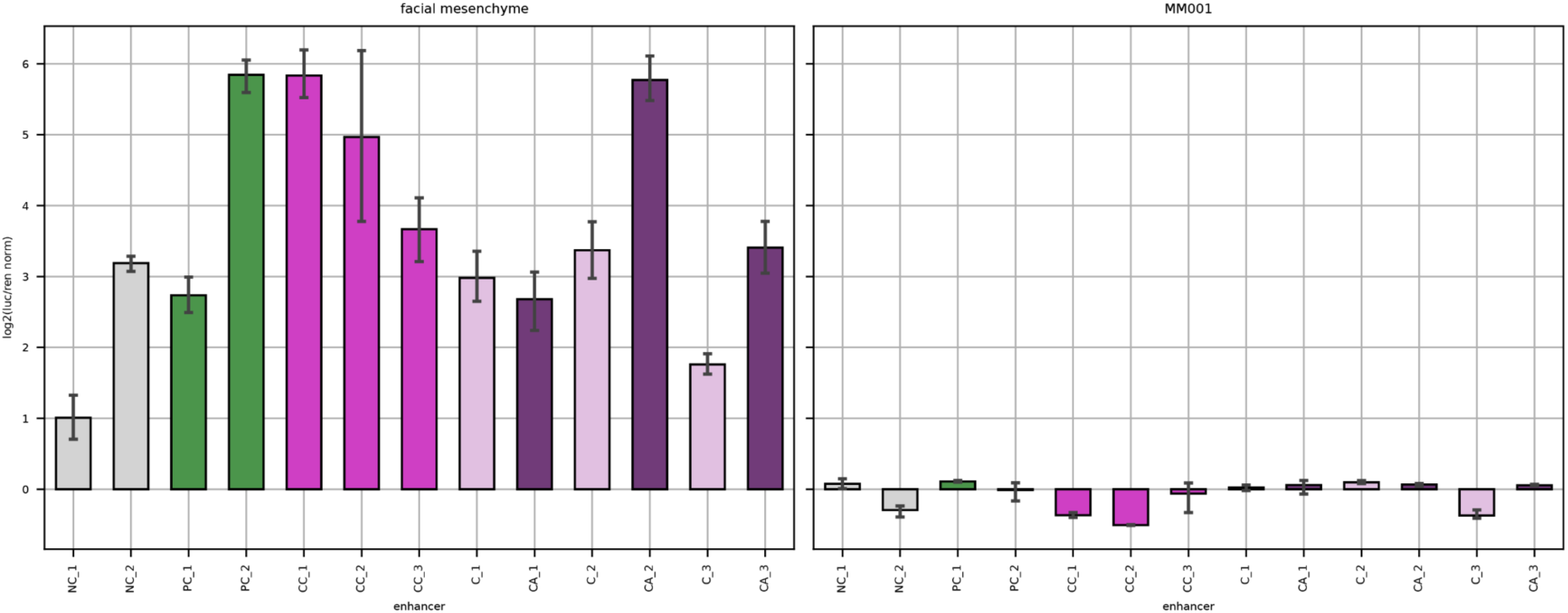
Luciferase reporter assay in facial mesenchyme cells and MM001 melanoma cell line. Luciferase reporter assay in facial mesenchyme cells (left; n=6 sequences) and MM001 melanoma cell line (right; n=3 sequences). NC: negative control (genomic region with low DeepNeuralTube prediction score); PC: positive control (genomic region with high DeepNeuralTube prediction score); CC: synthetic enhancers with two high-affinity coordinator instances; C: synthetic enhancers with one high-affinity coordinator instance. CA: synthetic enhancers with one high-affinity coordinator instance and five accessory instances.

**Figure S11.**
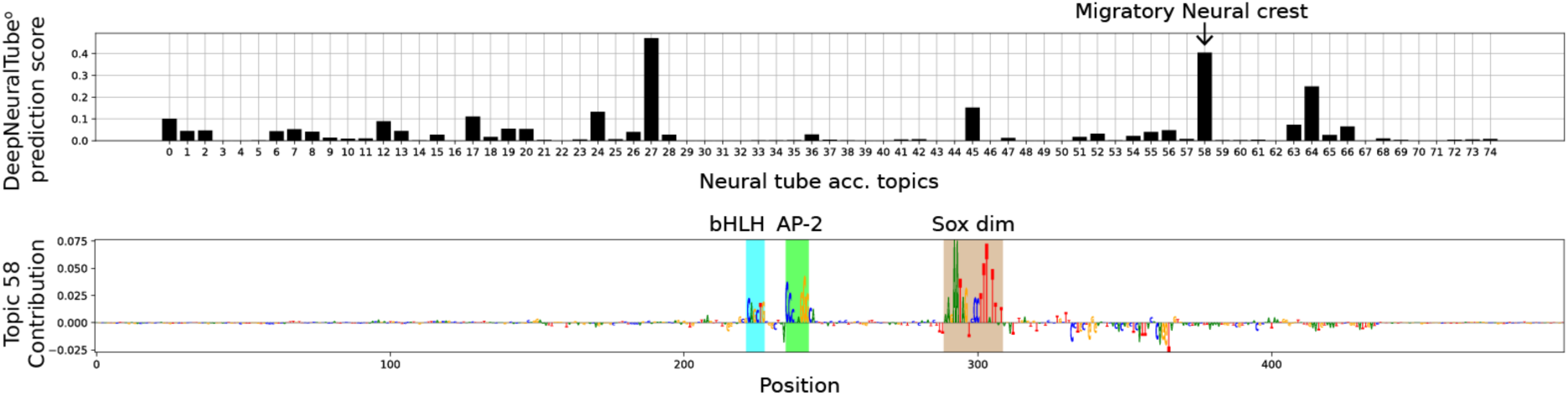
Prediction and contribution score for negative control enhancer NC2 predicts activity in migratory neural crest cells. DeepNeuralTube° prediction score (top) and contribution score for topic 58 (bottom) for negative control enhancer NC2.

## Supplementary tables

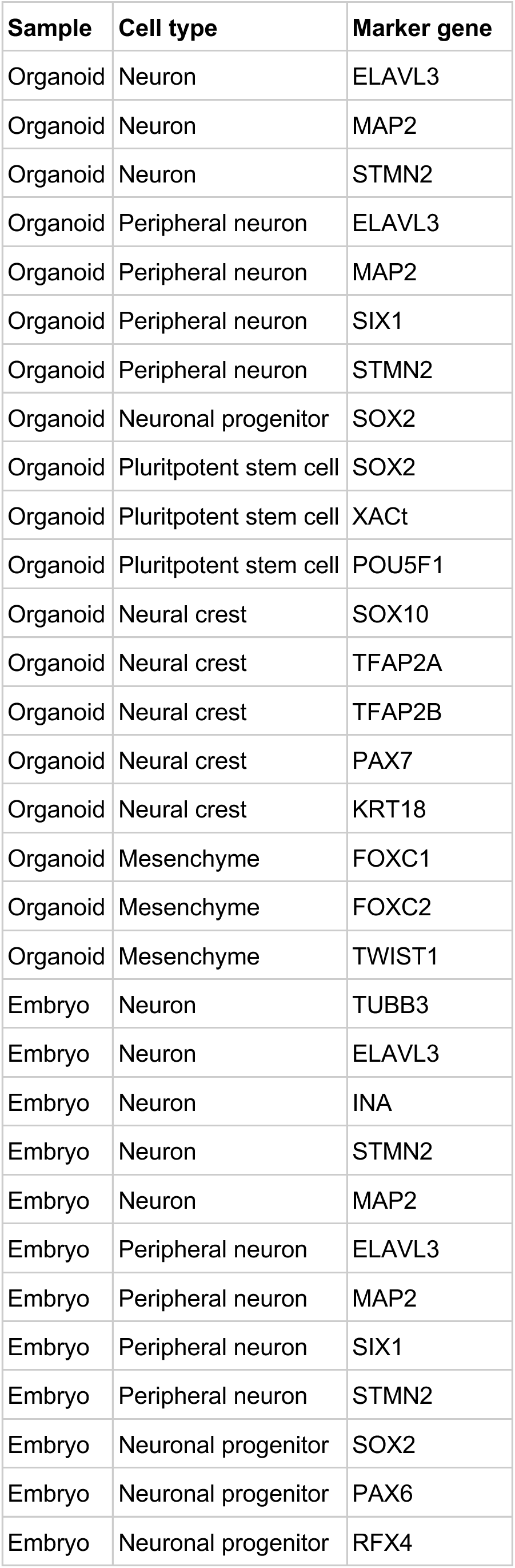
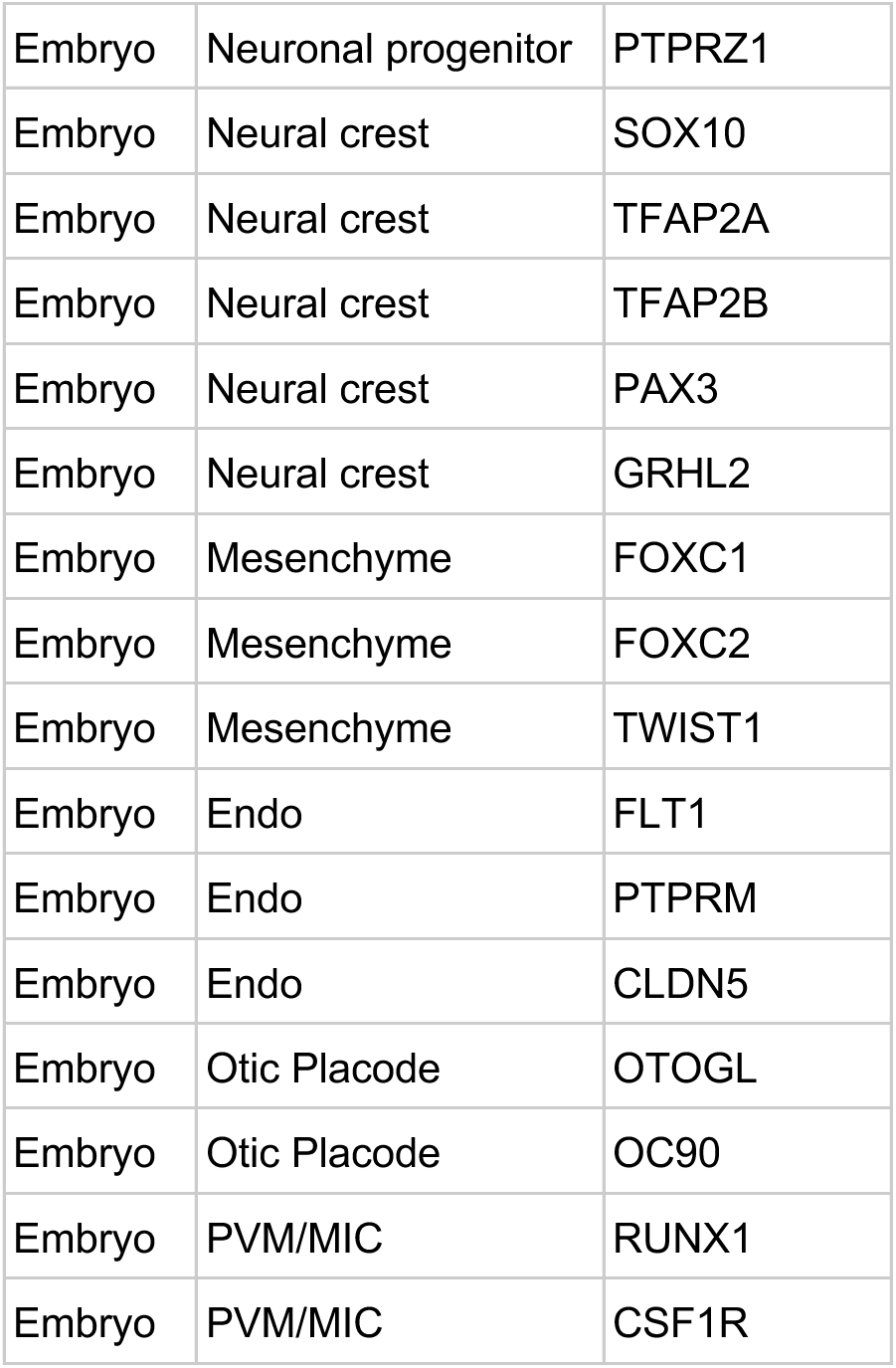
Table S1

## Notes

https://github.com/aertslab/TF-MInDi

https://github.com/SeppeDeWinter/NeuralTube

https://www.ncbi.nlm.nih.gov/geo/query/acc.cgi?acc=GSE295337

### Competing Interest Statement

The authors have declared no competing interest.

